# Biased replay of aversive and uncertain outcomes underlies irrational decision making

**DOI:** 10.1101/2025.01.17.633533

**Authors:** Tricia X. F. Seow, Jessica McFadyen, Raymond J. Dolan, Tobias U. Hauser

**Affiliations:** Max Planck UCL Centre for Computational Psychiatry and Ageing Research, University College London; London, United Kingdom; Functional Imaging Laboratory, Department of Imaging Neuroscience, University College London; London, United Kingdom; State Key Laboratory of Cognitive Neuroscience and Learning, IDG/McGovern Institute for Brain Research, Beijing Normal University; Beijing, China; Department of Psychiatry and Psychotherapy, Faculty of Medicine, University of Tübingen; Tübingen, Germany; German Centre for Mental Health (DZPG); Tübingen, Germany

## Abstract

Humans often make irrational decisions when facing uncertain or aversive future events despite careful deliberation. How we make choices, including irrational ones^1,2^, has been the object of extensive study both behaviourally and neurally^3–6^ and is the focus of influential behavioural economic theories^7–9^. Yet, little is known about how these (irrational) decisions are carved out in the brain. Here, using magnetoencephalography (MEG), we show that the construction and outcome evaluation of irrational decisions involves rapid, sequential state reactivation, or “replay.”^10^. During deliberation, we show that forward replay is biased towards choice options with more negative and uncertain outcomes, with this bias further amplified immediately preceding irrational choice. Likewise, post-decision evaluation relates to replay in a choice-dependent manner. Following irrational choices, relief-like signals were evident as stronger backward replay of worse counterfactual options, while after rational decisions, regret-like signals appeared as stronger backward replay of better counterfactual options. Together, these findings suggest that neural replay shapes both the formation and reflection of irrational decisions, and poise replay as a candidate mechanism underlying pervasive decision biases in humans.

## Introduction

Humans are prone to make decisions that defy logic or reason, for example when choosing to gamble despite knowledge that the odds are against them^11^. In behavioural economics, selecting options that fail to maximize beneficial consequences, as determined by weighting anticipated outcome value and uncertainty^7–9^ are characterised as irrational^1,12^. While prior studies identified several factors that explain why behaviour deviates from that predicted by a neoclassical utility model^2,13^, little is known regarding the neural antecedents of irrational preferences^14^. Thus while studies have identified brain activity that correlates with key decision variables^3–6^, a mechanistic account for processes underlying irrational decisions has remained elusive. One suggestion is from sequential sampling models of decision-making^15^, where variability in memory samples relate to (irrational) decisions^16^. An ensuing variability in representational contents should, in principle, be susceptible to measurement by neural replay, a phenomenon that involves a rapid sequential reactivation of representational states^17^.

Here, we address a hypothesis that neural replay mediates the construction of irrational decisions. Replay is proposed as a mechanism supporting spatial navigation^18,19^, where spatial trajectories play out in a forward manner during navigational planning, or backward when consolidating an outcome^18,20^. Replay events are implicated in cognitive domains that extend beyond spatial processing^21,22^ where developments in magnetoencephalography (MEG) data analysis^10^ have enabled characterisation of replay-like mechanisms in humans during risky approach/avoidance^23^, learning^24–26^, inference^17,27,28^ and recall^29,30^. Furthermore, as humans universally experience emotions of regret or relief^31,32^ with respect to outcome knowledge of unchosen alternatives^33^ after choice, we also examined the role of replay in the evaluation of decisions, given its known role in credit assignment^27^ and memory consolidation^29,30,34^.

In this study, we focus on the question of whether irrational decisions are driven by neural replay. Using an adaptation of a canonical risky decision-making task, we found evidence for forward replay when deliberating choice options. Critically, there was stronger forward replay for options with more negative and uncertain outcomes, which were amplified prior to an irrational choice. We also found evidence for post-choice backward replay, where a backward replay of counterfactual options was modulated in a regret or relief-like manner depending on choice rationality. Together, our findings implicate replay in the construction and reflection of irrational decisions.

## Results

Participants (N=30) completed a two-alternative forced choice decision task involving varying levels of uncertainty and outcome valence. Prior to this, participants were instructed that there were three (option) paths, each leading to one of three possible outcome types (reward (money), neutral, uncomfortable electric shock) (**Fig. 1A**). Each of these paths consisted of three images (“states”), with each outcome type also represented by an image. In the decision task, participants deliberated between choosing a probabilistic (probabilistically transitions to one of two paths; “gamble”) or a deterministic (transitions to one path; “certain”) option (**Fig. 1B**). On every trial, the transition probability, outcome type, and outcome value varied for each path (**Fig. 1C**). By using acquired knowledge of each path sequences and outcomes, participants could evaluate the expected value (EV, the multiplication of transition probability with its outcome value^7^) of each path/choice option (**Fig. 1D**)) using individually calibrated values of reward versus shock (**Supplementary Note 1**, **Supplementary Fig. 1**). A choice was considered rational if participants chose the option with the higher EV^7^.

**Fig. 1.**
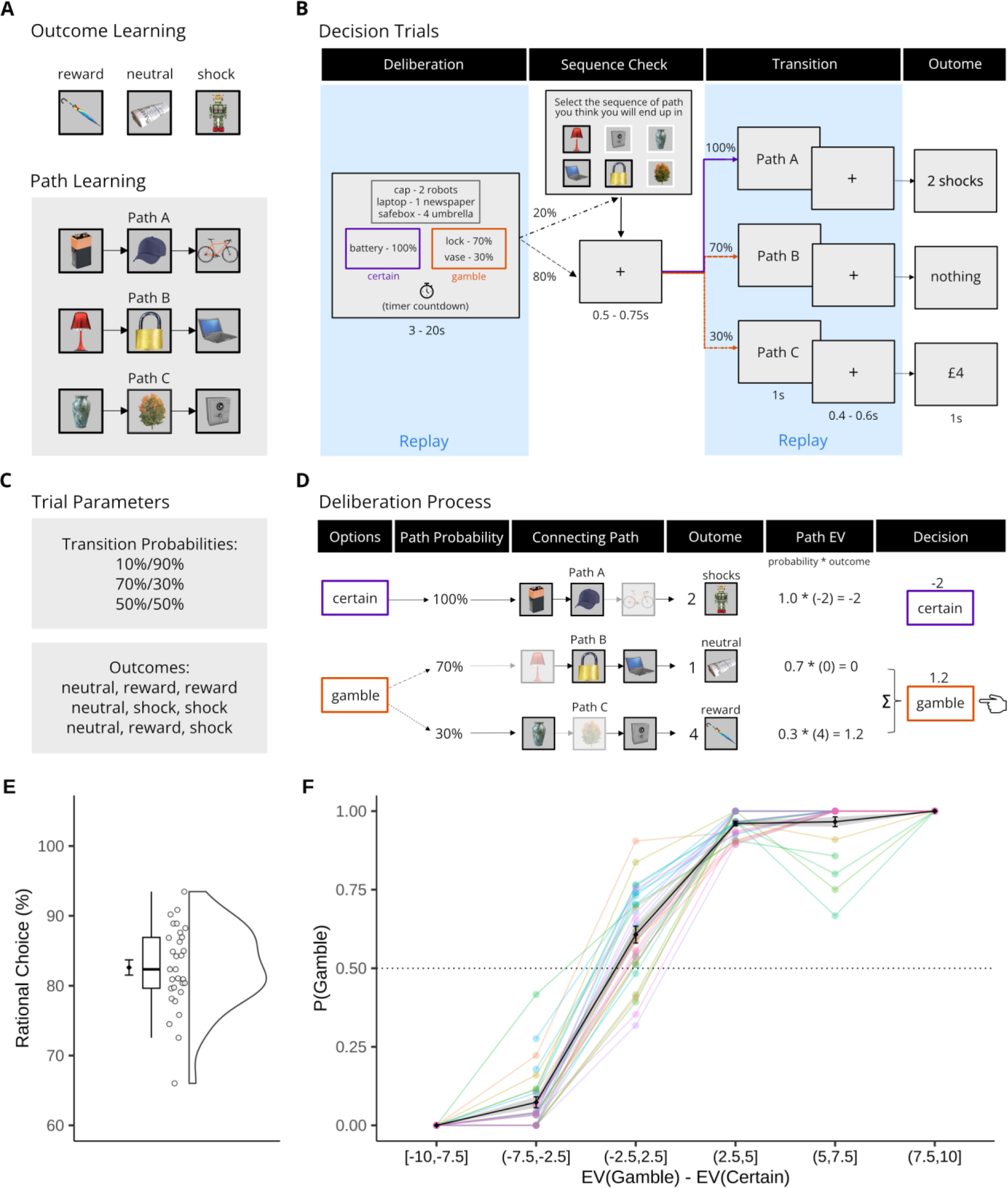
Decision task paradigm and behaviour. **(A)** Participants learnt the associations of images linked to a possible outcome (reward, neutral, shock) as well as three path sequences (path A, B, C) consisting of three images (“states”; e.g., S1-S2-S3 in path A). All participants showed good memory for this path information after a single training iteration (M=91.67%, SD=12.95%). **(B)** On each decision trial, participants were presented with cues. They deliberated between a probabilistic (gamble) or deterministic (certain) choice leading to one of three paths and their unique outcome. Once a choice was made, participants could encounter (20% of the time) a sequence check screen where they were tasked to select the state sequence of the path they expected to experience given their choice (score used as a co-variate in analyses). Participants were highly accurate in reporting the order of path states over the course of the task (M=92.63%, SD=6.72%). Thereafter, participants transitioned to a path (only path identity was shown, no states) and its outcome. The sections in blue indicate the time (deliberation and post-choice) where replay activity was quantified for further analyses. **(C)** The probabilities of the gamble option, outcome type combinations and their magnitude (1-5 shocks; monetary rewards matched in subjective value, c.f. **Methods**) for the three paths, varied across trials. The combination of probability and outcome type also varied (i.e., each path may lead to shock, nothing, or reward). See **Supplementary Note 3** and **Supplementary Fig. 3** for the decision trial schedules. **(D)** The cues for each trial in (**B**) encouraged deliberative planning. The first or second state from each path was linked to a transition probability, while the second or third state was linked to an outcome state (type) and value. Probability and outcome were always linked to two different states of a path. Pooling this information, participants could construct learned path sequences and link each path’s transition probability with its outcome value for informed deliberation of their choices. Making rational choices requires calculation of each path’s expected value (EV; probability*outcome value), with subsequent selection of the option with the higher EV. For illustration purposes, outcome values in (**C**) and (**D**) do not reflect individually calibrated reward to shock values (**Supplementary Note 1**, **Supplementary Fig. 1**). **(E)** Rational choice is defined on the basis of choosing the option with a highest EV. Participants made rational choices majority of the time, demonstrating that they perform the task well, albeit with individual differences. **(F)** Participants made certain or gamble choices based on comparison of path EVs. Brackets indicate the inclusion, while parentheses indication exclusion, of the value next to it. For (**E**) to (**F**), thin lines and data points indicate individual participants.

### Probability and outcome guide choice

Using EV as an index of rational choice, participants made significantly more rational (than irrational) choices (M=82.60%, SD=5.94%; t(29)=30.05, p<0.001, 95% CI=[60.77 69.65]) (**Fig. 1E-F**), demonstrating they understood the task adequately. Nonetheless, no single participant was entirely rational, with individual components of EV—options’ outcomes and probabilities—separately influencing the decision process (**Supplementary Note 2**, **Supplementary Fig. 2**).

### Significant replay during deliberation

To probe the content of what participants pondered whilst deciding, and how this related to the likelihood of an irrational choice, we concurrently recorded MEG data. In particular, we focused on neural replay of option paths during decision deliberation (3 to 20s; **Fig. 1B**). We first constructed multivariate state classifiers trained on visually-evoked whole-brain MEG activity for each unique state image (i.e., S1-S12), as derived from a pre-task functional localiser (**Fig. 2A-B**; c.f. **Methods**). We then applied these state classifiers to MEG deliberation time data to estimate reactivation of option path states and used a temporally delayed linear modelling (TDLM) framework^10^ to assess evidence for temporally-ordered reactivation (replay or “sequenceness”) of option path states (via valid state transition pairs of the paths e.g., S1-S2, S2-S3, S1-S3 in Path A) across different time intervals (10 to 600ms, in 10ms steps). Sequenceness was estimated for both forward (e.g., S1→S2, S2→S3, S1→S3) and backward (e.g., S3→S2, S2→S1, S3→S1) directions.

**Fig. 2.**
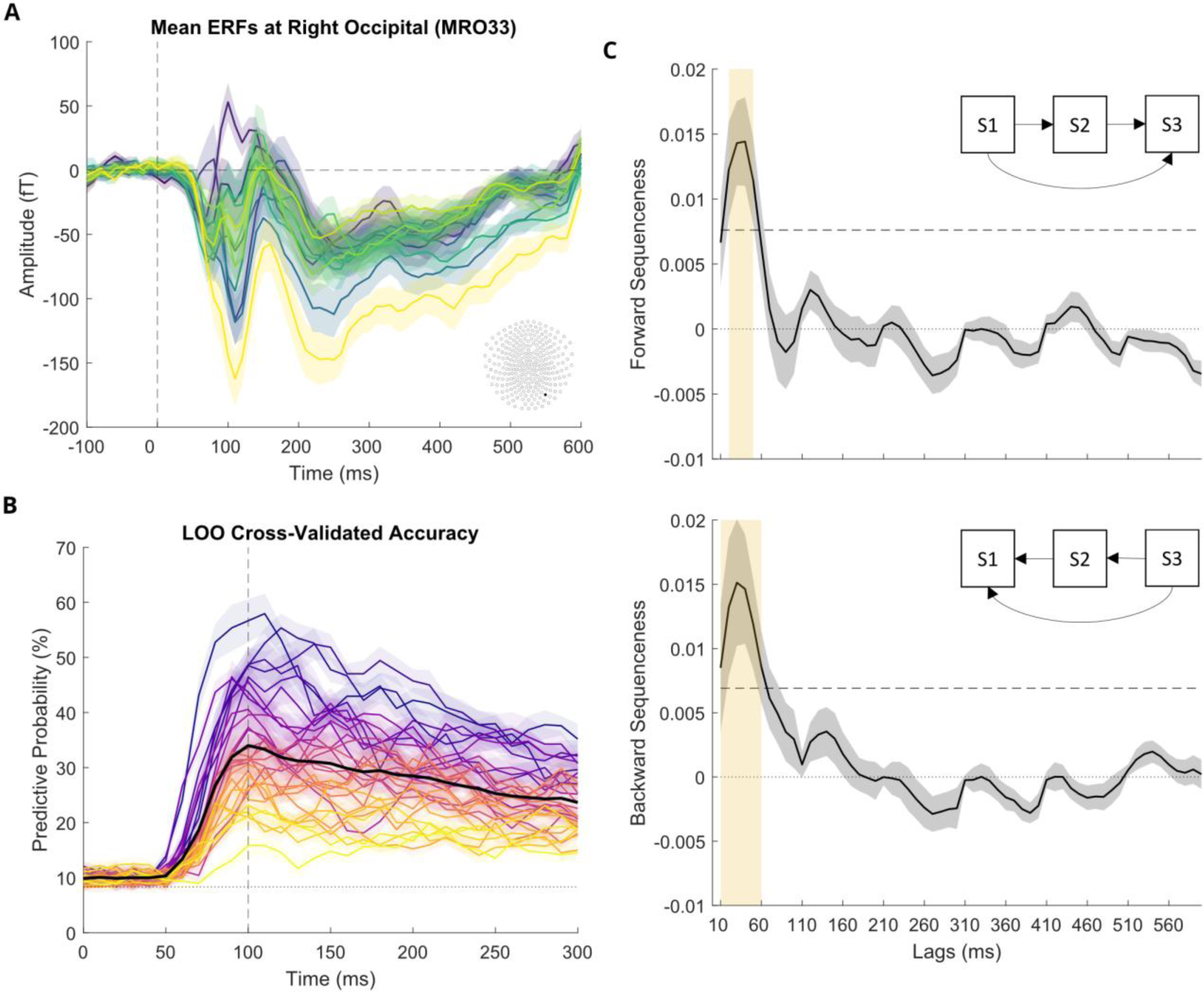
State classification and sequenceness (replay) quantification. **(A)** Participants partook in a functional localiser task that captured the visually-evoked event-related fields (ERF) of unique images (or “states”, S1-S12). Each individual colour represents the grand-average ERF for each unique state at a right occipital sensor (black dot on the sensor map) across participants. **(B)** For each participant and each state, a set of beta weights per sensor (classifier) was created based on stimuli-related activity from the functional localiser task. In a leave-one-out (LOO) cross-validation, we examined the probability (accuracy) of each state classifier in predicting its own state compared to all other possible eleven states. Across participants, classifiers trained at 100ms from onset of the stimuli showed the highest average accuracy overall. Each individual colour represents the predictive probability across all states for each participant. **(C)** Using temporally delayed linear modelling (TDLM)^10^, we estimated sequential state-to-state reactivation for a series of interval lags (10ms, 10ms to 600ms) in a forwards and backwards direction (c.f. **Methods**). Figures indicate the sequenceness estimate (or ‘replay’, indicated by the bold black line) averaged across all valid state transitions (e.g., S1-S2, S2-S3, S1-S3 for path A, given that path A was construed by states S1-S2-S3). This is the degree to which these state pair reactivations corresponded to the option paths’ sequential state structure (forward: S1→S2, S2→S3, S1→S3, and backward: S3→S2, S2→S1, S3→S1) for all option paths, across all participants. The significance threshold (95^th^ percentile of 199 null permutations) is indicated by the horizontal dashed line. Significant forward sequenceness was observed at intervals (lags) of 20-50ms, peaking at a 40ms interval, while significant backward sequenceness was observed at lag intervals of 10-60ms, peaking at a 30ms interval, as indicated by the yellow shaded rectangle. Shaded areas are the standard errors of the mean (SEM) of each state activity across participants in (**A**), the SEM of each participant classifier accuracy across states in (**B**), and the SEM of sequenceness across participants in (**C**).

During deliberation, we observed maximal sequenceness estimates, i.e., highest sequential replay (average of all valid state pair transitions) of (all three) option paths, at a 40ms state-to-state lag interval in forward (significant intervals: 20-50ms, non-parametric permutation test, c.f. **Methods**), and 30ms interval in backward (significant intervals: 10-60ms), direction (**Fig. 2C**). These lag intervals are consistent with a similar rapid temporal compression seen in prior human replay studies^17,23,29^.

### Forward replay during deliberation

Given significant forward and backward sequenceness during deliberation, we next probed whether replay carried decision-relevant information using linear mixed effect models. Specifically, we asked whether path replay (average sequenceness of all valid transitions for each path) was influenced by its outcome value and transition probability. We predicted that trial-by-trial sequenceness strength of each option path (for significant [forward: 20-50ms; backward: 10-60ms] intervals) would be modulated by outcome value or probability, controlling for potential confounders and co-variates (c.f. **Methods**). We found that prospective outcome influenced option path forward replay strength, with stronger replay for paths with more negative outcomes (β=-0.0008, SE=0.0003, p=0.008) (**Fig. 3A**). This effect was also evident when we probed forward replay strength between outcome types. Compared to reward paths, both neutral (β=0.002, SE=0.0008, p=0.01) and aversive (β=0.002, SE=0.0007, p=0.002) paths had higher replay strength. In contrast, we did not find any consistent backward replay effects in relation to outcomes (**Supplementary Note 5**).

**Fig. 3.**
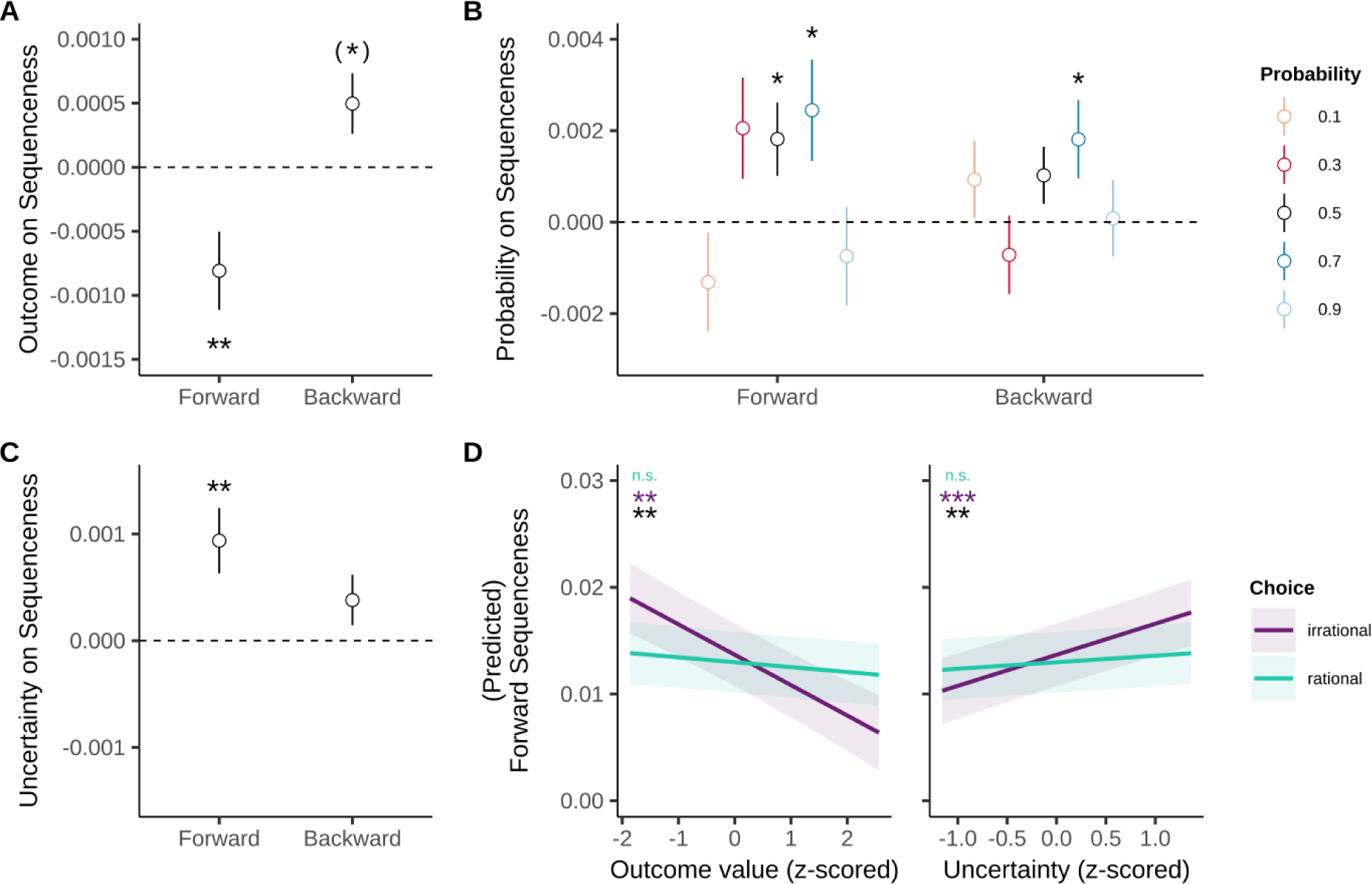
Forward, but not backward, replay is linked to path outcome and uncertainty, which predates irrational (versus rational) choice. **(A)** Forward sequenceness is linked to outcomes of the option paths, where positive option paths elicit less replay than aversive (and neutral) option paths; an association not seen for backwards replay. **(B)** In comparison to certain (100%) paths, uncertain (50% and/or 70%) paths had stronger forward sequenceness. **(C)** Probabilities recoded into uncertainty (100%: certain, 90/10% = a little uncertain, 70%30% = somewhat uncertain, 50%/50% = most uncertain) showed paths with high uncertainty exhibited stronger forward, but not backward, sequenceness strength. For (**A-C**), Y-axes are beta estimates from the linear models predicting sequenceness strength. Error bars indicate ±1 SEM. **(D)** Before an irrational choice, forward sequenceness was stronger for paths with more aversive outcomes and higher uncertainty transitions. Shaded areas are standard deviations. Significance indicated as teal = rational choice model, purple = irrational choice model, black = interaction effect with choice rationality. p^(^*^)^<0.05 but not a consistent effect (**Supplementary Note 5**), p*<0.05, p**<0.01, p***<0.001.

We next investigated whether forward replay was modulated by path probability and found that replay modulation by probability reflected its quadratic counterpart, namely uncertainty. Thus, forward sequenceness was stronger for more uncertain path transitions, such that compared to certain paths, both 70% (β=0.002, SE=0.001, p=0.03) and 50% (β=0.002, SE=0.0008, p=0.02) path transitions showed an increasing strength in forward replay (**Fig. 3B**). As uncertainty effects followed an inverted U-shape relationship of path probability, we recoded probability into uncertainty (50% (very uncertain) 70%/30% (uncertain), 90%/10% (somewhat uncertain), 100% (certain)). Again, we found that stronger forward sequenceness tracked the uncertainty of choice options (β=0.0009, SE=0.0003, p=0.002) (**Fig. 3C**), an effect not seen for backward replay (β=0.0004, SE=0.0002, p=0.11).

Lastly, in a model that included both decision variables, we found evidence for independent effects of outcome and uncertainty on forward sequenceness strength (outcome: β=-0.0008, SE=0.0003, p=0.006; uncertainty: β=0.001, SE=0.0003, p=0.002) suggesting that neural representations of decisional information relates to the uncertainty of options rather than raw probabilities. This contrasts with behavioural theories that posit irrational decisions stem from a skew of probability^35^, but is consistent with uncertainty being a strong driving factor in relation to subjective preferences^36^.

### Biased replay precedes irrational choice

Next, we probed the relationship between path replay and choice rationality, classifying each choice on the basis of having chosen the option with the higher EV (rational), or not (irrational). Before participants made irrational choices, we saw enhanced forward replay of aversive options (**Fig. 3D**; β=0.002, SE=0.0009, p=0.005; interaction between outcome and rational/irrational choice, c.f. **Methods**). More specifically, preceding an irrational choice there was enhanced replay of paths that lead to aversive outcomes (β=-0.003, SE=0.0008, p=0.001), with no evidence for such an effect when preceding rational choice (β=-0.0005, SE=0.0003, p=0.17). Thus, there was evidence for augmented deliberation-related replay of states leading to negative outcomes preceding irrational choice.

We also ascertained whether there was an effect of uncertainty and found forward replay was again tightly linked to irrational choice (**Fig. 3D**; interaction: β=-0.002, SE=0.0009, p=0.009). Thus, participants were more likely to make irrational decisions when there was enhanced replay of uncertain paths (β=0.003, SE=0.0008, p<0.001), but not so for rational choice (β=0.0006, SE=0.0003, p=0.06).

We found no effects of outcome value (p=0.42) and uncertainty (p=0.32) on backward sequenceness, nor any interaction with choice rationality (ps>0.66). This suggests that the exaggeration of value and uncertainty prior to irrational decision-making lay in forward, not backward, replay. While here we focused on outcome and uncertainty effects on replay that preceded choice, we nonetheless examined if choice rationality could be predicted from the replay of the three individual option paths. We found that deprioritised replay of the best choice option together with enhanced replay of the worst option predicted irrational choice (**Supplementary Note 6**). We also found that the addition of replay over just behaviour determinants increased the likelihood of predicting choice (**Supplementary Note 7**).

Finally, when testing whether outcome value or uncertainty effects on replay might be driven by choice difficulty, we found no effect (ps>0.29) nor any interaction with rational choice (ps>0.20) (**Supplementary Fig. 5**). To ensure replay effects were not driven by choice difficulty, in all analyses we include trial difficulty regressors as co-variates, ensuring we capture preference-related, rather than difficulty-related, effects.

### Backward replay after choice

Next we tested whether, akin to its proposed role in credit assignment^27^ and memory consolidation^29,30,34^, replay was also present during post-choice reflection, and whether it carried choice-related information. Here we focused on a timepoint when participants first learned about the consequence of their choice; immediately following a transition to the option path identity leading to outcome receipt (**Fig. 1B**). Similar to replay during deliberation, we found significant forward sequenceness at 40-50ms state-to-state lag interval, and backward sequenceness at 20-50ms interval, with both having peaks at 40ms interval (**Fig. 4A**) at this time.

**Fig. 4.**
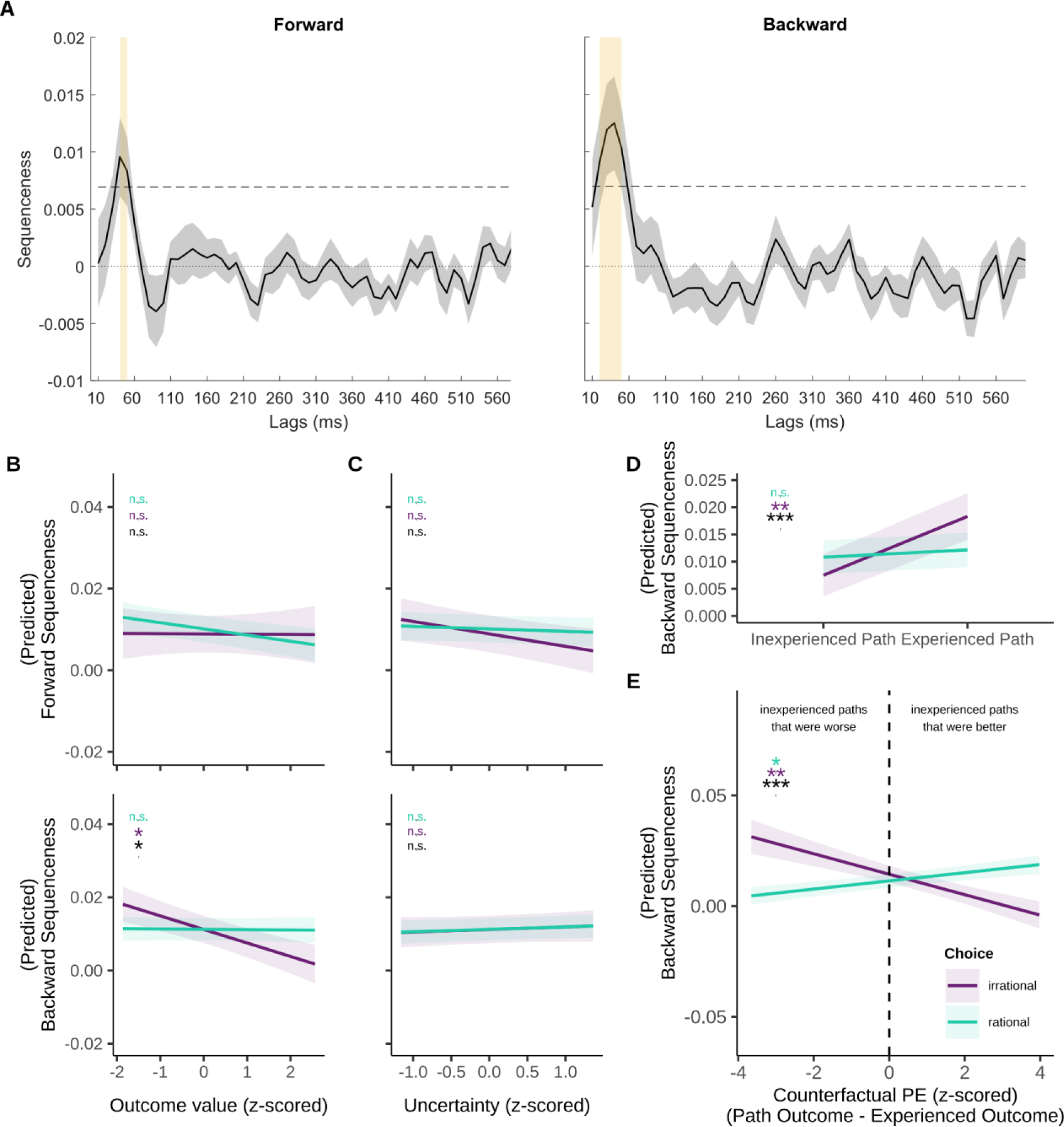
Replay at transition onset after choice. **(A)** Significant sequenceness post choice, both in forward (40-50ms intervals, peak at 40ms interval) and backward (20-50ms intervals, peak at 40ms interval) directions, are indicated by the yellow shaded areas. Dashed lines indicate significance threshold determined by 199 null permutations. **(B)** Backward replay after choice reflects outcome value. The directionality of sequenceness associated with outcome value switched after choice with stronger backward sequenceness seen for paths with more negative outcomes. **(C)** Absence of uncertainty effect on option path replay after choice. Transition uncertainty was resolved once participants were shown which path they have attained. Post choice, neither forward or backward sequenceness were associated to the transition uncertainty of the path. **(D)** Replay of attained paths (experienced) versus forgone paths (inexperienced). Participants replayed the paths they were transitioned to more strongly than the paths they were not, and this was particularly so after irrational choices. **(E)** Replay after choice is sensitive to counterfactual prediction error of option paths. At the transition screen, participants could compare the outcome value of the attained path (experienced) with the foregone paths’ value (inexperienced). For each trial, we estimated the difference between each path’s outcome and experienced path’s outcome, termed a counterfactual prediction error (PE). Experienced paths had a PE=0, and better alternative inexperienced paths had a PE>0 while worse alternative inexperienced paths had a PE<0. We found that more aversive inexperienced paths after irrational choice, together with better inexperienced paths after rational choice, were associated with stronger backward replay. This echoes the missed-out outcomes after bad choices (relief) or missed out better outcomes after good choices (regret). Shaded areas in (**A**) are SEM, and in (**B**) and (**C**) are SDs. n.s.>0 05, p*<0.05. Significance in teal = rational choice model, purple = irrational choice model, black = interaction effect with choice rationality.

Based on a proposal that replay reflects what is currently task relevant^29,37^, we would expect path replay relates to the yet to be presented outcome value (but where participants know which outcome they will receive), and not to path transition uncertainty/probability which is resolved fully at this path transition stage. In contrast to the pattern seen during deliberation, we found post-choice forward path sequenceness (40-50ms intervals) was no longer linked to outcome value (p=0.98), path uncertainty (p=0.18), or their interaction with choice rationality (ps>0.32) (**Fig. 4B-C**). Instead, we found evidence for backward sequenceness (20-50ms intervals) of paths linked to outcome value alone (β=-0.004, SE=0.002, p=0.02) (**Fig. 4B**) and where now resolved path uncertainty no longer impacted backwards replay (p=0.66; **Fig. 4C**).

We then saw that backward replay strength for aversive option paths was choice-dependent (interaction: β=0.004, SE=0.002, p=0.03) such that, following irrational choices, there was enhanced replay of aversive option paths (β=-0.003, SE=0.002, p=0.03), but this was not seen after rational choices (β=-0.00005, SE=0.0006, p=0.94). Moreover, the path leading to the to be received outcome was replayed more strongly than the other two foregone option paths (β=0.01, SE=0.003, p=0.001), particularly so after irrational choice (interaction: β=-0.009, SE=0.004, p=0.009) (**Fig. 4D**). This enhanced replay of path states leading to aversive outcomes thus appears to reflect an enhanced focus on bad outcomes consequent upon a poor choice.

### Counterfactual reflection after choice

A stronger replay for aversive outcomes after irrational choices suggests that participants were concerned with consequences of their decisions, a key basis for credit assignment and learning. Given replay has a role in both^24,27,29^, we hypothesised that replay of the option paths would be sensitive to a comparison of forgone possible outcomes to an actual received outcome. Conditional on whether participants made a good or bad choice, this would reflect a relief or regret signal respectively. On this basis, we estimated a counterfactual prediction error (PE) quantifying the difference between alternative paths (inexperienced paths) outcomes versus the outcome participants actually received (experienced path). Here, inexperienced paths would have a positive PE if these led to a better outcome than the one experienced (regret) and a negative PE if that path led to a worse outcome than experienced (relief).

We then tested whether this counterfactual PE, and its interaction with choice rationality, predicted backward sequenceness strength of the option paths. We found that a modulation of backward sequenceness in relation to counterfactual PE was dependent on choice rationality (interaction: β=0.006, SE=0.002, p<0.001) (**Fig. 4E**). In separate models that focused on either rational or irrational trials, paths with lower PEs (i.e., worse alternative paths than the experienced path) had stronger backward sequenceness after irrational choice (β=-0.005, SE=0.002, p=0.004), but weaker backward sequenceness after rational choices (β=0.002, SE=0.0007, p=0.01). This indicates that backward sequenceness signalled relief after irrational choice (increased replay for worse alternative paths; i.e., “I luckily did not get worse outcomes”), and regret after rational choice (increased replay for better alternatives; i.e., “I could have gotten better outcomes”).

We also saw that counterfactual PEs were negatively associated with backward replay (β=-0.005, SE=0.002, p=0.003) above choice-dependent effects. This indicated that inexperienced options with worse prospects expressed stronger backward sequenceness overall, reflecting a sense of general optimism (i.e., avoided these outcomes).

Finally, we highlight that backward replay strength was better explained by counterfactual PE value as opposed to outcome value (PE vs. outcome model: ΔAIC=-9.95, ΔBIC=-9.94). None of these effects were evident for forward replay (ps>0.63). We also note that other conceptualisations of counterfactual PE showed a similar effect (**Supplementary Fig. 6**). Overall, our findings suggest that post-choice backward replay reflects the retrospective evaluation of a decision, akin to relief and regret in a choice-dependent manner.

## Discussion

Despite the benefits of deliberation, humans are disposed to make choices that deviate from rationality^1,12,38^. Economic theories have highlighted that potential choice outcomes and their likelihood^8,9^, along with their brain representations^3–6^, as key tenets of decision-making. However, how irrational choices are constructed and evaluated, and whether neural representations of decision features influence the pondering of a choice has been underexplored. In this study, we show that specific expressions of human neural replay is involved in the construction of irrational preferences and the subsequent shaping of post decision evaluations.

During deliberation, forward replay of option paths was modulated by both outcome value and expectancy. We observed greater forward replay for aversive and uncertain option paths, an effect that was significantly amplified prior to irrational choices. This increased replay sensitivity coupled to an enhanced replay of the worst option paths preceding irrational choices suggests that replay prioritisation of un-preferred paths determines subsequent choice^23,39^. Interestingly, we note that the opposite (replay of an aversive path predicted its avoidance, not approach) pattern is described in rodent replay^40^; albeit the latter was proposed to reflect recall of stored trajectories and not the planning process per se. We also highlight that replay patterns reflected uncertainty rather than raw probabilities, in line with prior proposals of an uncertainty-organised brain^41^. This contrasts with traditional behavioural suggestions that humans use skewed probability representations in decisions leading to irrational behaviour that manifest as underestimating rare, or overestimating common, events^35^.

Our findings suggest that a distorted representation of key decision variables drives choice irrationality^1^. Viewed from the context of sequential sampling models of decision-making^15^, this implies that instead of variability in memory samples underlying choice, the replay patterns we describe reflect decision option weights. This extends findings showing that neural activation is preferentially related to computations involved in weighing value versus probability for risk^42^. Importantly, it implies that replay reflects more than just memory stability^30,40^, given that outcomes and contingencies are trial-independent and presented numerically on-screen for the entire deliberation time in our task design.

Models of replay propose that it acts as a general learning mechanism^34,43,44^, which would assume importance during retrospective evaluative processes. In line with this, we show that post choice backward replay of inexperienced option paths reflect a comparison between a foregone and experienced outcomes. Notably, we found a general positive post-choice reflection evident in stronger replay for alternative paths leading to worse outcomes (i.e., “at least I avoided that”). We also found a choice dependent replay prioritisation of “what might have been” with stronger replay for worse alternative options after irrational choice, and stronger replay for better alternative options after a rational choice—echoing counterfactual retrospective evaluations that induces relief (‘could have gotten worse’) and regret (‘could have gotten better’) respectively^45^. As the one-shot choices in the current task do not require state-action values updating for sequential decisions^24,34^, these replay patterns would appear to instantiate computations that relate to subjective reflection as opposed to learning^46^.

We found a marked change in decision-related replay directionality—forward replay during deliberation and backward replay during post-choice—without any instructions/requirements to utilise the paths in a particular direction. This aligns with theories about replay directionality serving a specific computational goal^34,47^, such as forward replay for prospective cognition (e.g., planning of future states^24,25^) and reverse replay for retrospective evaluations (e.g., credit assignment^27^ and memory consolidation^29,30^). Thus, our findings extend and integrate previous findings related to replay directionality and task demands^17,23–26,29,30^, suggesting a rapid replay directional change may be fundamental to the dynamics of cognitive processes.

It is noteworthy that deliberative biases are amplified in mental health disorders, exemplified in debilitating indecision and reassurance-seeking behaviors^48,49^. In this context, our supplemental analyses showed deliberative choice-rationality dependent outcome replay biases in individuals with high obsessive-compulsive or anxious-worry traits (**Supplementary Fig. 8**). This finding aligns with proposals of aberrant replay underlying psychiatric symptoms^23,26,50^, and suggest biased replay as a candidate process for quantifying neurocognition across the psychopathological spectrum.

In summary, we demonstrate that deliberative and evaluative mechanisms of choice are strongly tethered to neural replay, opening new avenues to probe more pervasive deliberative and reflection biases, particularly in psychiatric illnesses with ruminative dysfunctions.

## Methods

### Participants

We recruited N=30 healthy volunteers from the UCL Psychology subject pool (SONA). All participants were between 18-55 years, had no history of psychiatric (assessed with the Structured Clinical Interview for DSM-V (SCID)^51^) or neurological disease/injury, had no history of certain medical conditions (cardiac disorder, uncontrolled hyperthyroidism, severe hypertension or significant skin conditions), were not taking any psychiatric medication, were fluent in English, had normal/corrected-to-normal vision, and met all criteria for MEG scanning. Participants provided informed consent for both the online practice and in-person imaging session. They were compensated £8.50/hour, plus up to £5 depending on their performance on various components of the study. The final study sample consisted of participants 18-46 (M=27.17, SD=8.09) years old and 21 (70%) females.

### Procedure

Study procedures were conducted in accordance with and approved by the London Westminster NHS Research Ethics Committee (15/LO/1361). Participants were first remotely assessed in a screening interview for their psychiatric history (SCID) and criteria for MEG scanning. If participants were deemed eligible, they performed the behavioural task practice and a battery of questionnaires online. Subsequently, they attended the in-person imaging MEG session that included the following modules:

### Shock Voltage Calibration

We used mild electric shocks as the aversive outcome in the experimental task. Shocks were stimulated by a Digitimer high voltage constant current stimulator model DS7A, with pulse duration set at 1000μs. Delivery of the shocks were controlled by a custom MATLAB script, where 1 shock unit was defined as a quick train of 5 stimulations with 16ms intervals. We calibrated individualised shock voltage levels for each participant, following a standardised procedure where participants were given 1 unit of incrementally higher voltage shock repeatedly to the back of their hand. After every increase, they were asked to rate its intensity on a scale from 1 to 10, where 10 was the highest level that they would be willing to tolerate. Intensity was slowly increased until a level of 10 was reached. This procedure was repeated 3 times. We used 80% of the final recorded level 10 voltage intensity as the aversive outcome in the experimental task, same as other protocols^25^.

### Reward-Shock Staircase

Following the shock voltage calibration, the participants completed a task intended to balance the value of shock and reward for the estimation of expected value by estimating a reward-shock ratio (R:S). This ratio (ranged from 0.05-1.15 (M=0.60, SD=0.37)) determined how much money was rated equal to 1 shock unit, and was subsequently used in the main experimental task to transform the original reward value. In other words, if a participant had a R:S of 0.8, they equated £0.80 to 1 shock. In the decision task, we would then assume that +0.80 (reward) and -1 (shock) were adjusted in parity. A random trial from the reward-shock staircase procedure and the main experimental task determined the number of shock units participants acquired, which was delivered at the end of the session. See **Supplementary Note 1** for details of the reward-shock calibration procedure.

### Functional Localiser

Participants were then set up to scan with MEG. They first performed a functional localiser task. On each trial, an image was presented on screen for 500ms, followed by two words on the left and right. One of the words was the correct label for the image shown while the other word was randomly selected from a pool of invalid labels. Participants pressed either the left or right button of a 2x4-button response pad to indicate the correct label. They had a maximum of 100ms to make a response, or the trial would be skipped. After making a response, the chosen word was indicated by a white box surrounding the word for 250ms. A fixation cross was then shown for a randomly jittered inter-trial interval between 500-700ms before the start of the next trial. Participants completed 600 trials split over 4 blocks, where 12 unique images (cat, bicycle, hourglass, bowtie, car, backpack, zebra, toothbrush, baby, lamp, house, and cupcake; utilised in other replay studies^23^) were presented 50 times each in random order. The participants’ accuracies in this task were high (M=97.55%, SD=1.44%).

### Experimental Task

The experimental task was programmed in MATLAB R2019b with Psychtoolbox v3.0.18. In the decision paradigm (**Fig. 1**), participants played the role of an astronaut taking one of two space shuttles from a central command centre to get to three different spaceship types. Each spaceship type (path) contained a sequence of three rooms (path sequence). Thus, there were three path sequences: spaceship A contained rooms S1, S2 and S3, spaceship B contained rooms S4, S5, and S6 while spaceship C contained rooms S7, S8 and S9. All path sequences would also lead to an outcome room, where the participant would either gain money (S10), receive shocks (S11), or is neutral/nothing (S12). Each room was represented by a unique image from the image array utilised in the functional localiser task, randomly allocated per participant. In other words, each path sequence contained 3 images, while each outcome (reward, shock, neutral) was represented by 1 image.

### Outcome & Path Learning

Participants first learned the associated images to the possible outcome types (reward/shock/neutral) (**Fig. 1A**) before being tested on this knowledge. Subsequently, they learned the associated images of the rooms with the spaceships (path sequences). Participants were presented with the sequence of the 3 room images for each spaceship twice, with each image presented for 2000ms with a fixation cross of 250ms in between. They then completed a quiz on the order of the path sequences where they were to select the image that represented a room in a particular spaceship (2-arm forced choice) and what room image came before/after that room (3-arm forced choice).

### Decision Paradigm

After completing outcome and path learning, participants then started on a tutorial for the main task. At the beginning of each trial, participants were presented with a cue screen (**Fig. 1B**). They were told that they had a choice of taking one of two space shuttles, where one shuttle would deterministically (100%) lead to one spaceship, and the other would probabilistically (50-50%, 70-30%, or 90-10%) lead to one of two other spaceships. The identity of these spaceships was indicated by either the first or second room image of each spaceship associated to its transition probability. Participants were also told each spaceship would lead to a certain outcome room (ranging 1-5 shocks, neutral, or 1-5 equivalent rewards), which was associated with either the second or third room of each spaceship. With these two pieces of information, participants were able to use the path sequences learnt beforehand to decipher which shuttle would take them to which spaceship and to predict what outcome they will gain from their choice. We assumed that the neutral outcome (“nothing”) had a value of 0, reward outcomes were positively valued (>0), and shock outcomes were negatively valued (<0). As such, participants were able to evaluate the expected value of choosing either the probabilistic (gamble) or deterministic (certain) shuttle option. Rationally, this evaluation would reflect an expected value calculation for both choices such that:

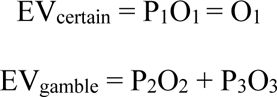

Where P is the probability of the path and O is the outcome, and the subscripts indices indicate each of the three different option paths. The decision to choose the deterministic option was considered correct (rational) if EV_certain_ ≥ EV_gamble_ and the decision to choose the probabilistic option was considered rational if EV_gamble_ ≥ EV_certain_. The number of trials with certain outcome combinations were pre-determined by 1 of 5 decision schedules randomly assigned to participants (**Supplementary Note 3**).

To prevent accidental presses and hasty responses, choices were only allowed after 3000ms. If a choice was made, the chosen option would be outlined in black for 300ms. If no response was made after 20000ms, the participant was penalised with the text ‘You lose £1! Please respond faster.’, shown for 1500ms.

For each block of trials, the first three trials were ‘forced choice’, meaning that participants were only allowed to choose the deterministic (100%) shuttle. They were then taken to a ‘sequence check screen’ where they were asked to indicate in sequence from a selection of 6 room images, the room order of the spaceship that they think they will end up in. Incorrect responses were determined as selecting a path sequence that does not exist (regardless of which path it reflects). Feedback on whether the participant has chosen a correct sequence (‘Correct order! No £ lost!’) or an incorrect sequence (‘No such order! You lose £1!’) will last for 1000ms. A randomly jittered fixation cross lasting 500-750ms was played before a screen indicating which spaceship they transitioned from their prior choice for 1000ms. This was followed by path sequence of the said spaceship, starting with a 250ms fixation and then one room image at a time for 750ms, with a 250ms fixation in between each image. A randomly jittered fixation cross lasting 400-600ms was shown before the final outcome image displayed for 2000ms.

For the subsequent trials of the block (‘free choice’), the sequence check screen only appeared 20% of the time, and no transition sequences (i.e., room images) was played. Instead, after the randomly jittered fixation (500-750ms) post-choice, a screen indicated which spaceship (path) they transitioned to for 1000ms. Then, a randomly jittered fixation cross lasting 400-600ms was shown before the final screen conveyed the outcome received in text for 1000ms. All trials had an inter-trial interval of a 1500ms fixation.

The reward and shock outcome values ranged from 1-5 in our construction of the decision trial schedules (**Supplementary Note 3**). Before the main task, the reward outcome values were transformed (multiplied) with the R:S (**Supplementary Note 1**) to balance the value of shocks and money for each individual. Furthermore, all outcome values were jittered with -0.1, 0 or +0.1 to reduce the identification of the outcome types without relying on learned images associations. This meant that if a participant had R:S=0.8 and the trial outcomes were determined as £2 and 2 shocks, the values seen on the cue screen would be £1.7 and 2.1 shocks (assuming +0.1 jitter for both, non-calibrated values). Before analysis, the reward outcomes values were transformed back with the R:S to reflect their true internal valuation of the reward that has equal parity as shocks (**Supplementary Fig. 1**). Thus, the reward value would be 2.125, and the shock value would be - 2.1 (calibrated values).

Before the start of the decision paradigm, participants played a tutorial of n=9 sample trials with sequence check after choice. Thereafter, they were required to take a final quiz on the associated images of the outcome rooms and the room sequences of the spaceships. In the decision paradigm, participants completed 9 blocks with MEG, where each block contained n=20 trials (3 forced choices, 17 free choices). In total, n=180 trials were performed. We note that 1 participant was only able to finish 7 blocks (140 trials) due to time constraints during their session.

Prior to the in-person MEG session, participants completed a shorter, online version of the experimental task. The online task was fully programmed with the JavaScript library React v.18.2.0 (https://react.dev/), developed in an app bootstrapped by Create React App (https://github.com/facebook/create-react-app), and hosted on Scalingo (https://scalingo.com/).

Shock outcomes were replaced by the prospect of losing money. Participants had to score >65% accuracy in the outcome learning, >75% in the path learning and 100% accuracy in the final knowledge quiz before they could play the decision task. The aim was to ensure that participants were capable of learning and performing the task for the MEG session. In total, participants played 6 blocks of 3 forced choice trials and 14 free choice trials each (n=102 trials) in the decision paradigm. Crucially, the images utilised in the online session were different from those shown in MEG session.

### Magnetoencephalography (MEG)

We recorded MEG data continuously at 600 samples/second with a 275-channel axial 586 gradiometer whole-head system (CTF Omega, VSM MedTech). 2 channels (MRC12, MLO42) were out of commission, leaving 273-channels for data collection. Participants were seated upright in the scanner and their head position was monitored by three head position indicator coils located at the nasion and left and right pre-auricular fiducial points. MEG triggers were recorded and retimed to a photodiode positioned behind the stimulus presentation screen that detected the onset of a flashing white/black stimulus (hidden from view) that was synchronised with event onsets. Participants’ eyeblinks were also recorded using an Eyelink eye-tracking system (SR Research). MEG data from the functional localiser and decision task were preprocessed with custom code written in MATLAB R2019b (https://www.mathworks.com/), using SPM12 (https://www.fil.ion.ucl.ac.uk/spm/software/spm12/) and Fieldtrip v. 20211102 (https://www.fieldtriptoolbox.org/).

### MEG Preprocessing

Data were first high-pass filtered at 0.1 Hz to reduce slow drift, and a notch filter for 50 Hz was applied to remove line frequency. The data were then downsampled to 100 Hz. Eyeblink correction was run semi-automatically with custom code based on Eyelink data. Independent component analysis was performed on the data, and components visually identified as eye movement and heartbeat artefacts were manually rejected. All subsequent analyses were performed directly on the filtered, cleaned MEG signal, in units of femtotesla. The data were then divided into different epochs using the trigger onsets and durations. For the functional localiser, epochs were created for the image onset (-100 to 600ms stimulus onset), which were baseline corrected with -100 to 0ms data. For the decision trials in the main task, epochs were created at deliberation time (-100ms cue screen onset to individual trial choice time) and at post-choice transition time (-100ms to 1400ms transition onset).

### Image classification

We first classified patterns of multivariate neural activity evoked by each image in the functional localiser task. For each stimulus (*i* ∈ {1:12}), we trained separate one-versus-rest L1 Lasso regularised logistic regression models using data from the functional localiser, from 0 to 600ms stimulus onset in 10ms time bins, excluding trials/epochs rejected for response inaccuracy or artefacts (M=47.70 trials per stimulus/participant, SD=1.42). Each model thus consisted of a trials x sensors (e.g., 600 x 273) data matrix and a binary trial image indictor vector (yes/no stimulus *i*). We ran these models with a range of 100 regularisation parameters (λ) sampled from a half-Cauchy distribution (γ = 0.05, range = 0.0001 to 1), producing a λ × sensors (100 × 273) matrix of slope coefficients, as well as a vector of intercept coefficients for each λ. These coefficients are the classifiers that discriminated sensor patterns of one stimulus versus all other stimuli.

To quantify classifier accuracy, we conducted a K-folds cross-validation procedure over every time bin where K was set to the minimum number of trials per stimulus for that participant. In each fold, one trial per stimulus were taken as the test set (n=12) while the remaining data was the training set. We then generated classifiers for each stimulus with the training set, and then tested if each classifier maximally predicted their correct stimulus from within test set (scored 1 or 0).

The average of scores across folds was taken as the accuracy of each classifier. For each participant, we selected λs that produced the highest mean accuracy across classifiers (M=0.0032, SD=0.00081). Overall average classification accuracy exceeded chance (8.33%) and peaked at 100ms (N=9/30).

### Replay Quantification

We first applied our classifiers to the MEG data during deliberation and post-choice time to estimate the degree to which the images (states) were sequentially reactivated in the brain by multiplying the spatiotemporal MEG data by each classifier’s beta estimates. Then, we used a 2-level Temporal Delayed Linear Modelling (TDLM) approach^10^ (https://github.com/YunzheLiu/TDLM) which quantifies the evidence for specific ordered state-to-state transitions with lagged cross-correlation, producing a “sequenceness” statistic at different time intervals (“lags”).

At the first-level, we evaluated each state-to-state reactivation pattern via assessing the average likelihood that stimulus *i* is followed by stimulus *j* after a time lag of *Δt*. We conducted multiple regressions for each path state’s reactivation time series (*i ∈* [1:9]), where the original, unshifted reactivation time series of state *i* (*X*_*i*_) is predicted by a time-lagged copy reactivation time series for another state *j* (*X*(*Δt*)_*j*_) with:

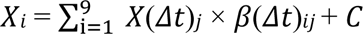

where *C* is a constant term and we used ordinary least-squares to derive the coefficient *β*(*Δt*)_*ij*_ that captures the unique influence of *X*_*i*_ on *X*(*Δt*)_*j*_. Thus, the first level coefficients form 9×9 empirical transition matrices, *B*(*Δt*), for each time lag. Separate models were estimated for each stimulus and each time lag *Δt* that ranged from 0 to 600ms in 10ms increments, where smaller intervals indicate more time-compressed reactivations.

At the second-level, we quantified the evidence for hypothesised ordered state-to-state transitions for each time lag. For the decision task, the key state-to-state transitions were S1→S2, S2→S3, S1→S3 (path A), S4→S5, S5→S6, S4→S6 (path B) and S7→S8, S8→S9, S7→S9 (path C). With the empirical transition matrices *B*(*Δt*) from the first-level regression, the evidence for the hypothesised transitions was then quantified by:

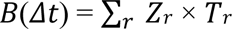

where *T*_*r*_ is the predictor transition matrix (state x state) for regressor *r*. We considered three predictor matrices (i.e., *r* ∈ {1:4}): (i) hypothesised forward (*T*_*F*_) transitions (e.g. S1→S2, S2→S3, S1→S3, etc.) set to 1 and all other transitions set to 0, (ii) hypothesised backward (*T*_*B*_) transitions (e.g. S3→S2, S3→S1, S2→S1, etc., the transpose of *T*_*F*_, (iii) an identity matrix of self-transitions (*T*_*auto*_) to control for autocorrelation, and (iv) a constant matrix (*T*_*const*_) that controls for general background neural dynamics. This enabled the estimation of *Z_r_*, which is the scalar regression coefficient quantifying the evidence for the hypothesised state-to-state transitions. Thus, *Z*_*F*_ and *Z*_*B*_ are evidence for forward and backward transitions, respectively.

We also computed a transition direction agnostic evidence measure *Z_C_* by using a summed *T*_*F*_ and *T*_*B*_ binary matrix for the second-level regression. To account for inter-subject variability in classification accuracy across training times and their relevance to replay, we followed a prior study procedure^23^ to elucidate the classifier training time per participant (90ms (N=8), 100ms (N=9), or 110ms (N=13); ±10ms the average maximum classifier accuracy training time) that produced the greatest absolute value of *Z_C_* across lags averaged across all transitions at deliberation time. These were then applied to quantify *Z*_*F*_ and *Z*_*B*_ sequenceness at deliberation time and post-choice time.

To determine the statistical significance of *Z* (“sequenceness”; averaged over the 9 valid transitions, all trials per participant), we used non-parametric permutation testing at the second-level. We generated 199 unique possible invalid versions of *T*_*F*_ and *T*_*B*_ (e.g., only included cross path transitions like S1→S4, S2→S5, etc.) and computed their (null) versions of *Z* to determine an empirical null distribution. Values of *Z* were deemed statistically significant (FWE<0.05) if they exceeded the significance threshold determined by computing the 95th percentile for *Z*_*F*_ and *Z*_*B*_ (one-sided test) of the maximum absolute value of each null distribution.

### Mixed-effects modelling

All replay analyses were conducted in R v.3.6.0 via RStudio v.1.2.1335 (http://cran.us.r-project.org). We utilised the t.test() function (stats package) for two-tailed paired t-tests and lmer() function (lmerTest package) to estimate mixed-effects models. Only free choice trials (n=153) of the MEG session were analysed.

A mixed-effects modelling approach allowed us to examine effects on replay on a trial-by-trial basis and compare conditions with unbalanced trial numbers.

To probe the influence of option components on its replay strength, we constructed a series of models that used sequenceness (forward or backward; Replay) as a linear dependent variable. The fixed effects co-variates included decision time (RT) and sequence memory score of the trial block (Seq Check), as greater RT naturally leads to a larger window to estimate higher sequencessness and sequence memory may affect sequencessness strength of the path options. As for random effects, we included the effects of per participant (Subject) as well as replay lag interval (Lags) as a nested random effect given that sequenceness was significant over multiple lags, all of which we included. Regressors were all z-scored.

To examine the relationship between replay strength with outcome value (Outcome) or transition uncertainty (Uncertainty), the models were:

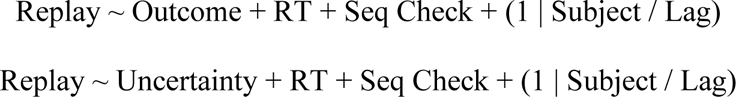

To examine the relationship between replay strength with outcome value (Outcome), transition uncertainty (Uncertainty) and its interaction effect with choice rationality (Rationality; yes or no), the model was:

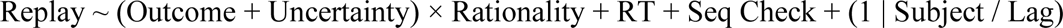

We added task difficulty regressors, where we adjusted for general trial difficulty (Choice Difficulty = abs(Gamble EV – Certain EV)) as well as the tendency for an easier gamble choice (Gamble Difficulty Bias = Gamble EV – Certain EV).

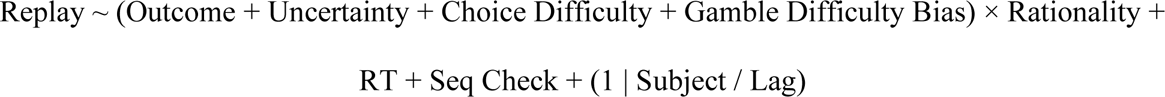

We estimated the counterfactual prediction error (PE) for every path, where each path’s PE was determined by the difference between its outcome, and the trial’s experienced outcome. The equation amounts to:

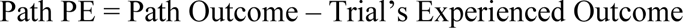

This meant that a path with PE=0 were paths that were ultimately received as the outcome within the trial. All other missed-out/inexperienced paths would have PE>0 or PE<0, where the former were inexperienced paths that had a better outcome, whilst the latter were inexperienced paths that had a worse outcome, then the one actually received. See **Supplementary Note 9** for other quantifications of PE.

To examine the relationship between post-choice path replay strength with outcome, experience of the path (i.e., if the transition led to the path identity or not), or counterfactual PE of the path and its interaction effect with choice rationality, the models were:

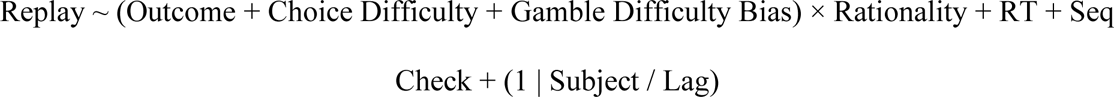

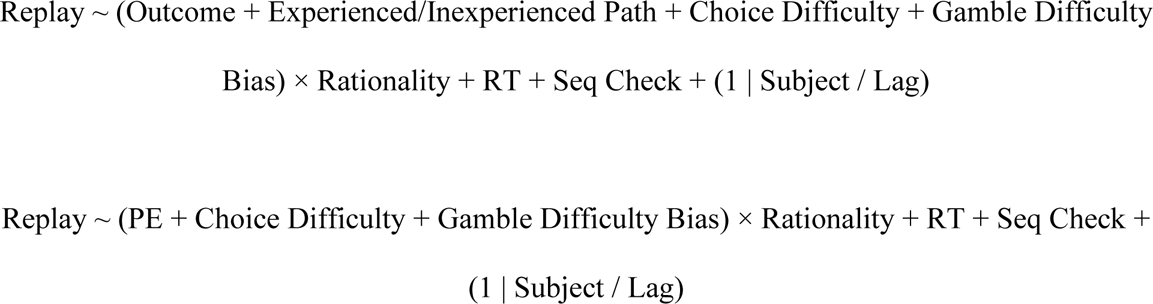

## End notes

## Acknowledgments

We thank Peter Dayan and Elliott Wimmer for their comments and discussion on the manuscript.

TXFS is a Sir Henry Wellcome Postdoctoral Fellow (224051/Z/21/Z) of the Wellcome Trust, based at the Max Planck UCL Centre for Computational Psychiatry and Ageing Research. JMcF was supported by a Wellcome Investigator Award (098362/Z/12/Z) to RJD. TUH is supported by a Sir Henry Dale Fellowship (211155/Z/18/Z; 211155/Z/18/B; 224051/Z/21) from Wellcome & The Royal Society and a grant from the Jacobs Foundation (2017-1261-04). The Wellcome Centre for Human Neuroimaging is supported by the Wellcome Trust (203147/Z/16/Z). This research was funded in whole, or in part, by the Wellcome Trust (211155/Z/18/Z).

The Max Planck UCL Centre for Computational Psychiatry and Ageing Research is a joint initiative supported by UCL and the Max Planck Society. For the purpose of Open Access, the author has applied a CC BY public copyright license to any Author Accepted Manuscript version arising from this submission. The funders had no role in study design, data collection and analysis, decision to publish, or preparation of the manuscript.

## Author contributions

TXFS and TUH conceptualized the study. JMcF and RJD contributed to the study design and methodology. TXFS coded and performed the experiment, analysed the data, and wrote the first manuscript draft with supervision from TUH. TXFS, JMcF, RJD and TUH reviewed and edited the manuscript.

## Data and materials availability

Data and the analysis code to generate the figures in this manuscript are available for download from an Open Science Framework repository (https://osf.io/sr74e/).

## Materials & Correspondence

Correspondence and requests for materials should be addressed to TXFS.

## Additional Information

Supplementary Information is available for this paper.

