## Supplementary information for "Biased replay of aversive and uncertain outcomes underlies irrational decision making"

Tricia X. F. Seow[1,2]*, Jessica McFadyen[1,2], Raymond J. Dolan[1,2,3] & Tobias U. Hauser[1,2,4,5]

[1]Max Planck UCL Centre for Computational Psychiatry and Ageing Research, University College London; London, United Kingdom

[2]Functional Imaging Laboratory, Department of Imaging Neuroscience, University College London; London, United Kingdom

[3]State Key Laboratory of Cognitive Neuroscience and Learning, IDG/McGovern Institute for Brain Research, Beijing Normal University; Beijing, China

[4]Department of Psychiatry and Psychotherapy, Faculty of Medicine, University of Tübingen; Tübingen, Germany

[5]German Centre for Mental Health (DZPG); Tübingen, Germany

### Supplementary Information Guide

#### 1. Supplementary Methods

| | |
|---|---|
| **Supplementary Method 1** | Factor analysis of questionnaire sub-scores |

#### 2. Supplementary Data

| | |
|---|---|
| **Supplementary Figure 1** | Estimating the balance of monetary value and shocks |
| **Supplementary Figure 2** | Decision behaviour |
| **Supplementary Figure 3** | Decision trial schedule example |
| **Supplementary Figure 4** | Cross validation of choice prediction without and with replay |
| **Supplementary Figure 5** | Choice difficulty effects on replay |
| **Supplementary Figure 6** | Counterfactual prediction error (PE) and replay after choice |
| **Supplementary Figure 7** | Individual differences in mental health symptoms |
| **Supplementary Figure 8** | Individual differences in mental health symptoms and replay |
| **Supplementary Figure 9** | Depression and reward to shock ratio |
| **Supplementary Table 1** | Predicting rational choice with replay |
| **Supplementary Table 2** | Factor analysis of mental health questionnaires |

#### 3. Supplementary Notes

| | |
|---|---|
| **Supplementary Note 1** | Calibrating rewards versus shocks |
| **Supplementary Note 2** | Decision behaviour |
| **Supplementary Note 3** | Decision trial schedules |
| **Supplementary Note 4** | Trial-to-trial independence of decisions |
| **Supplementary Note 5** | Inconsistent backward replay effects during deliberation time |
| **Supplementary Note 6** | Predicting rational choice with replay |
| **Supplementary Note 7** | Cross validation of choice prediction without and with replay |
| **Supplementary Note 8** | Choice difficulty effects on replay strength |
| **Supplementary Note 9** | Quantifying counterfactual prediction error after choice |
| **Supplementary Note 10** | Replay patterns in mental illness |

### 1. Supplementary Methods

**Supplementary Method 1: Factor analysis of questionnaire sub-scores**

Participants completed set of 6 questionnaires, assessing trait anxiety using the trait portion of the State-Trait Anxiety Inventory (STAI)[1], obsessive-compulsive disorder using the Obsessive-Compulsive Inventory - Revised (OCI-R)[2], impulsivity using the Barratt Impulsivity Scale (BIS-11)[3], depression using the Self-Rating Depression Scale (SDS)[4], obsessive beliefs using the Obsessive Beliefs Questionnaire - Short Version (OBQ-20)[5], worry using the Worry Domains Questionnaire - Short Form (WDQ-SF)[6].

We used a factor analysis (fa() from the psych package) with oblique rotation to reduce the dimensionality of 22 sub-scores from the 6 self-report questionnaires. We determined a three-factor structure with the Cattell-Nelson-Gorsuch scree test (nCng() from nFactors package)[7]. According to the top item loadings on each factor, we labelled them 'depressive-affect' (DA), 'obsessive-compulsive' and 'anxious-worry' (AW) (**Supplementary Fig. 7**; **Supplementary Table 2**).

### 2. Supplementary Data

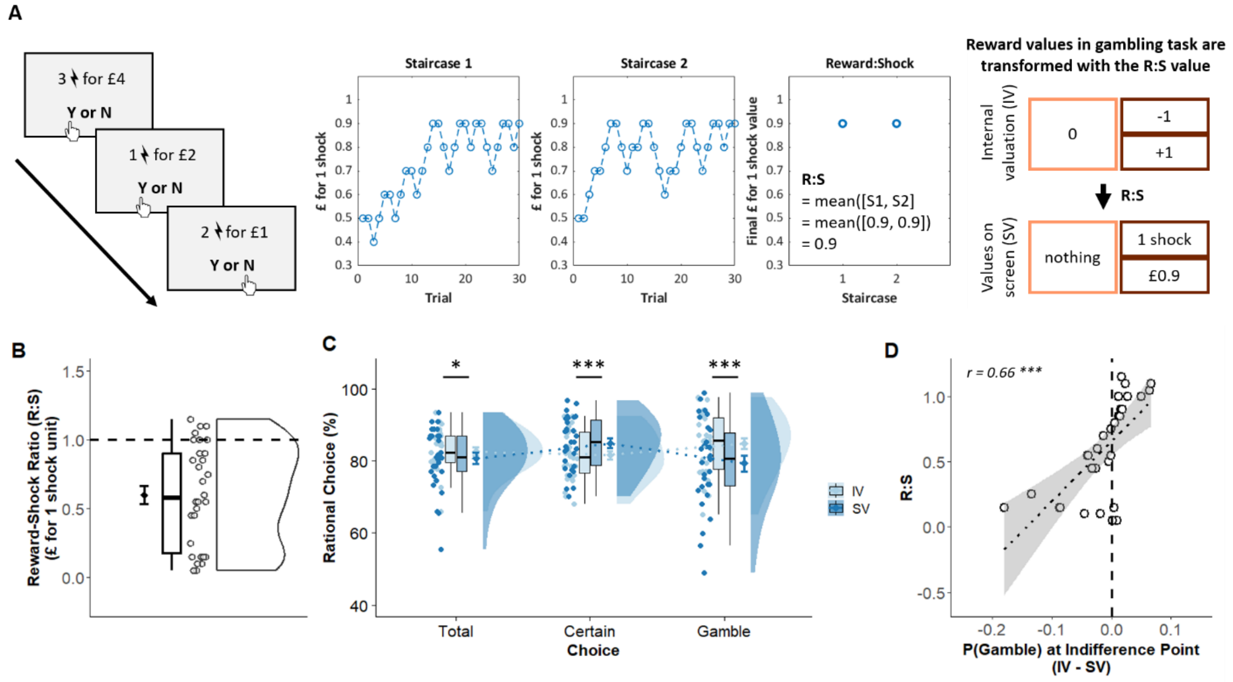


**Supplementary Figure 1. Estimating the balance of monetary value and shocks.**

(**A**) The reward to shock calibration paradigm consisted of a series of choices where participants decided whether to accept or reject various combinations of reward and shock units. Two underlying interleaved staircases determined the ratio between monetary value and shock units (reward to shock ratio; R:S), where the mean of the final values was equated to the amount of money a participant was willing to receive given 1 unit of shock. The R:S was used to transform the reward values in the trials of the main decision task, which ensured that the internal valuation (IV) of the options were comparable across participants, despite on-screen values (SV) being different. (**B**) Participants exhibited individual differences in the monetary valuation of 1 shock unit, the R:S. (**C**) Participants' choice behaviour were more rational overall when taking the IV values of the options, rather than the SV. (**D**) Whether participants were more risk-seeking or safe-seeking defined by their probability to gamble when choice options expected values were equal, differed when taking IV versus SV, which corresponded to their R:S value. The higher the R:S value, indicating a participant who was highly sensitive to shocks, the more risk-seeking they presented in IV than SV.

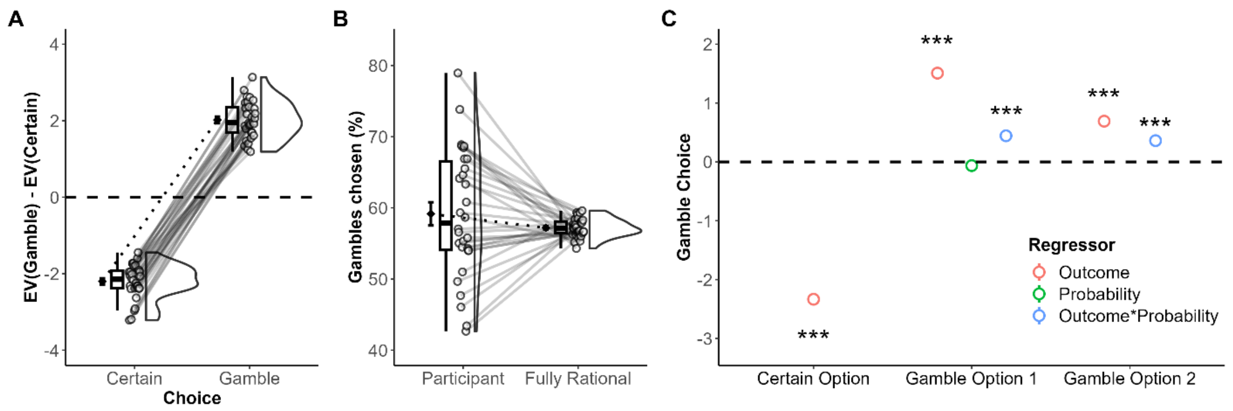


**Supplementary Figure 2. Decision behaviour.**

(**A**) The expected value (EV = probability*outcome value) of the gamble option (additive of both gamble paths EV) was higher than the certain option when participants chose to gamble and vice versa. (**B**) Participants selected more gamble than certain choices on average, in alignment to if choices were made in a fully rational manner. (**C**) We probed the separate contribution of outcome and probability on choice, showing that lower outcome values of the certain option together with higher outcome values of the gamble options increased gamble choices. Probability modulated the outcome effect on choice, where more likely gamble paths which had high outcome values predicted gamble choices even better.

For (**A**) to (**C**), thin lines and data points indicate individual participants. p***<0.001

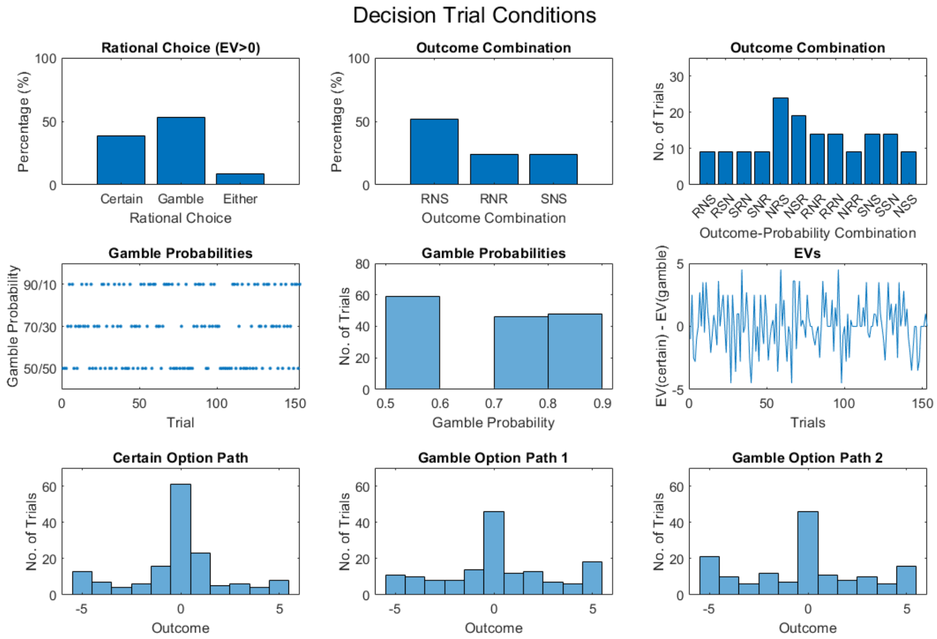


**Supplementary Figure 3. Decision trial schedule example.**

Participants completed one of five randomly assigned decision trial schedules determining the free choice (n=153) trials' option outcomes and probabilities.

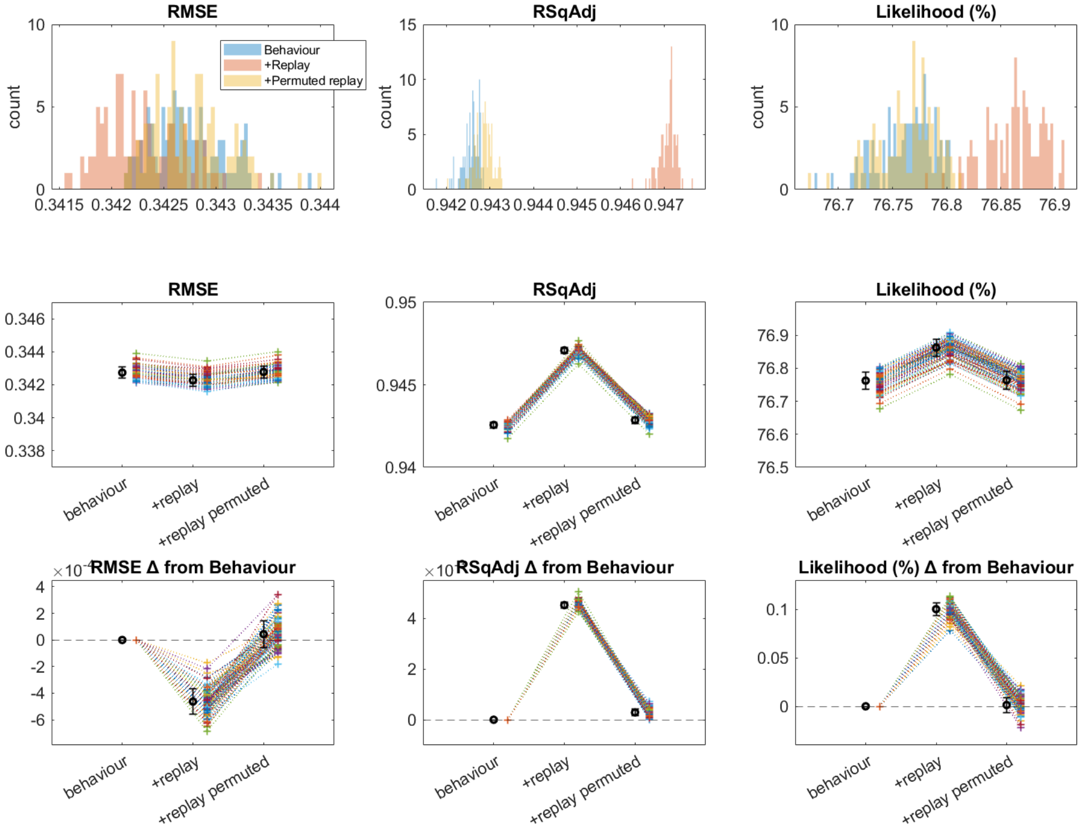


**Supplementary Figure 4. Cross validation of choice prediction without and with replay.**

We examined three different models to predict choice (certain/gamble) with a 5-fold cross validation procedure with 100 repetitions. The models included (i) a behaviour only model, (ii) a model including replay of the paths, and (iii) a model with replay that was randomly shuffled (permuted) across trials, which constituted a null distribution for comparison with (ii). We assessed root mean square error (RMSE), adjusted $R^2$ (RSqAdj) and likelihood of the models. Top row demonstrates the distributions of the model fit across 100 repetitions. In the middle and last rows, each coloured cross indicates the model fit from one repetition, averaged across the 5 folds, while the black marker indicates the mean and standard deviation of the model fits across the repetitions. The model including replay was the winning model, with the lowest RMSE and highest adjusted $R^2$ and likelihood. See **Supplementary Note 7** for details of the cross validation.

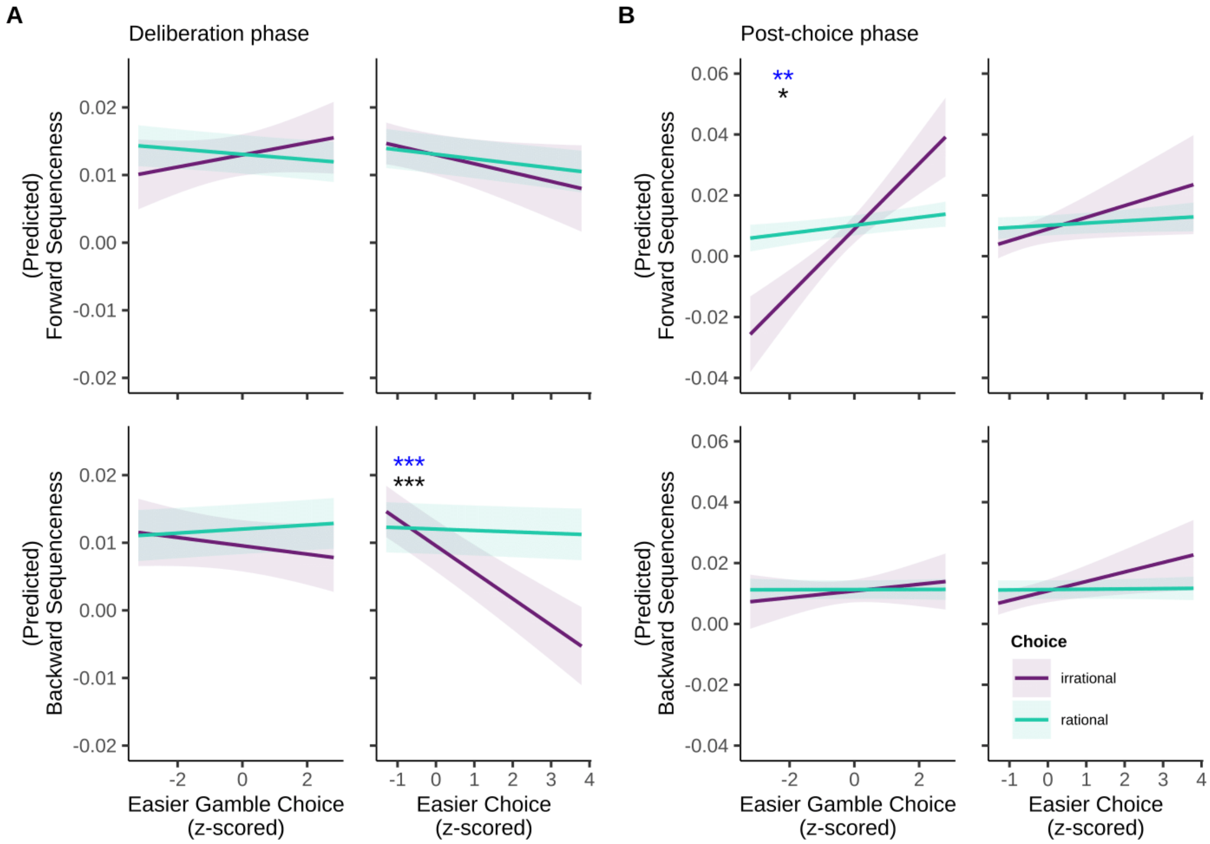


**Supplementary Figure 5. Choice difficulty effects on replay.**

(**A**) During deliberation phase, lower backward replay was observed for easier trials, particularly prior to irrational choice. (**B**) During post-choice phase, lower forward replay was observed for trials that were easier to make gamble choices, particularly after irrational choice.

p*<0.05, p**<0.01, p***<0.001. Blue = main effect, black = interaction effect.

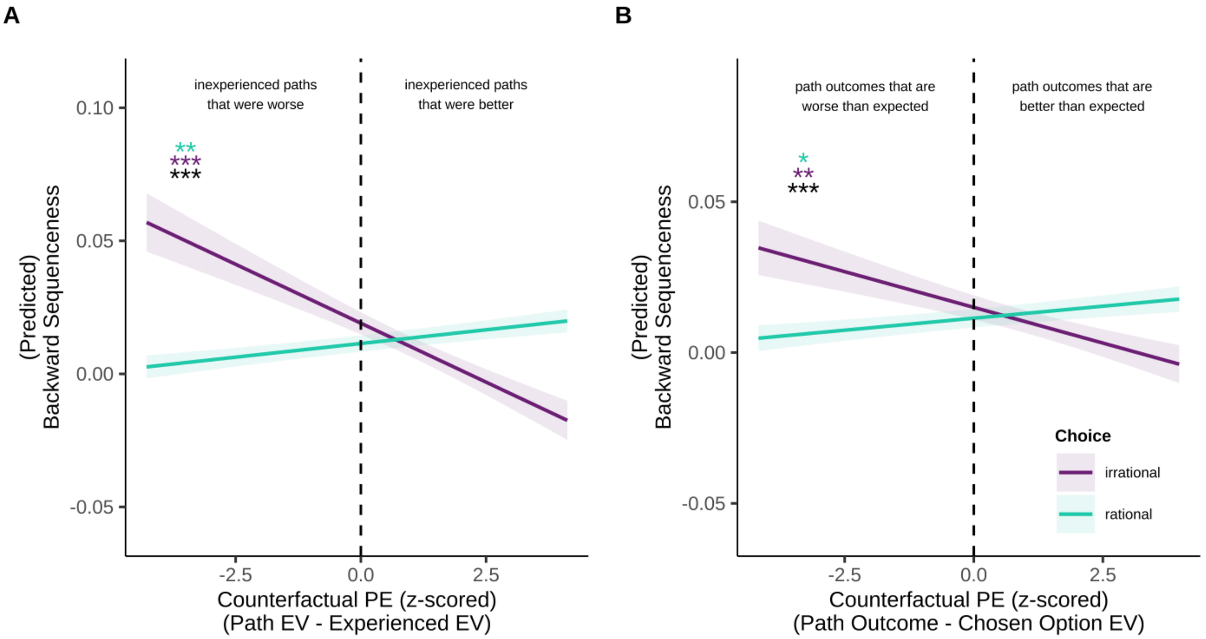


**Supplementary Figure 6. Counterfactual prediction error (PE) and replay after choice.**

(**A**) We estimated the PE as the difference between each path's EV and experienced path's EV. Therefore, experienced paths had a PE=0, and better alternative inexperienced paths had a PE>0 while worse alternative inexperienced paths had a PE<0. We found that the former had stronger backward sequenceness strength after rational choice, whilst the latter had stronger backward sequenceness strength after irrational choice. (**B**) Here, PE is the difference between each path's EV and the chosen option's EV. Therefore, option paths that had better outcomes than expected of the choice made had PE>0, while option paths with worse outcomes than expected of the choice made had PE<0. We found that the former had stronger backward sequenceness strength after rational choice, whilst the latter had stronger backward sequenceness strength after irrational choice.

Shaded areas are SDs. p*<0.05, p**<0.01, p***<0.001; teal = rational choice mode, purple = irrational choice model, black = interaction effect.

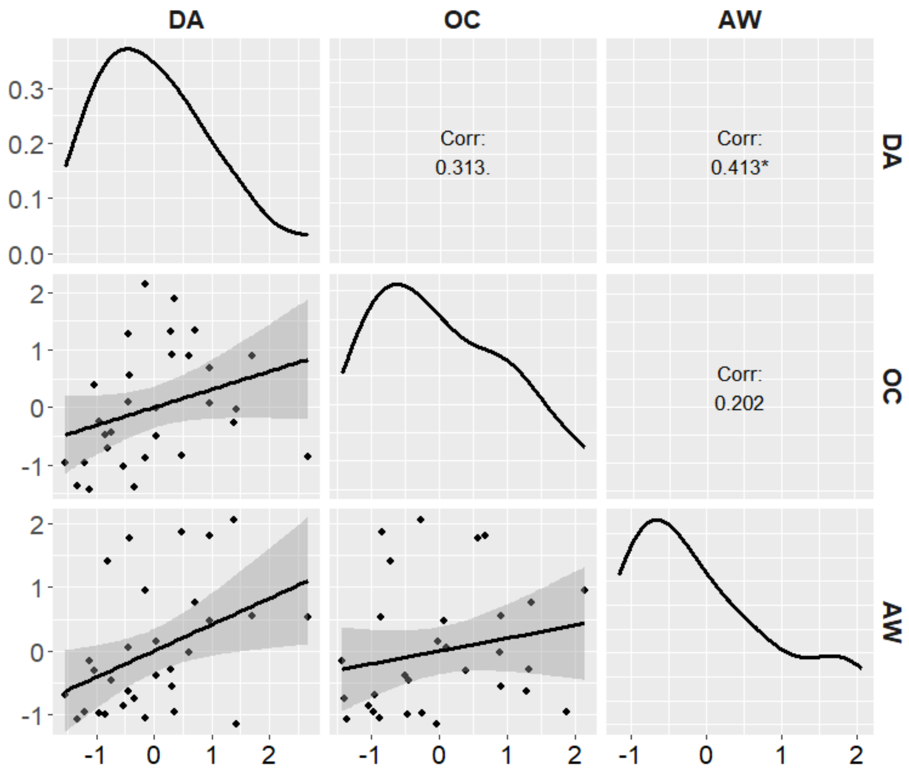


**Supplementary Figure 7. Individual differences in mental health symptoms.**

We characterised participants according to mental health factors of anxious-worry (AW), depressive-affect (DA) and obsessive-compulsive (OC) dimensions elucidated from a factor analysis of 22 sub-scores from 6 self-report questionnaires. Each individual datapoint is the factor score of each participant.

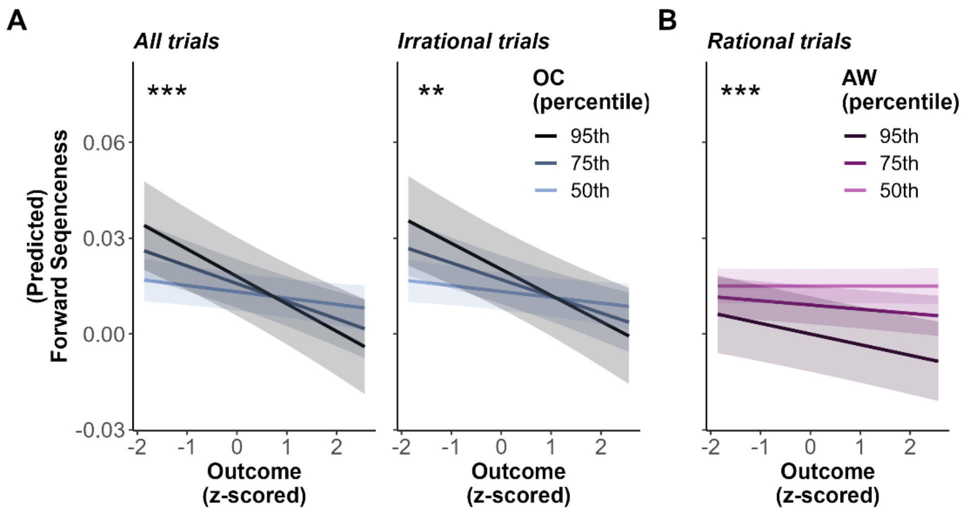


**Supplementary Figure 8. Individual differences in mental health symptoms and replay.**

(**A**) Before choice, an increased sensitivity of forward sequenceness to outcome value was seen within individuals high in obsessive-compulsive symptoms, with even stronger replay for aversive outcome paths before irrational choices. (**B**) In contrast, individuals high in anxious-worry had stronger replay for more negative paths before a rational choice.

Shaded areas indicate 95% confidence intervals. p**<0.01, p***<0.001.

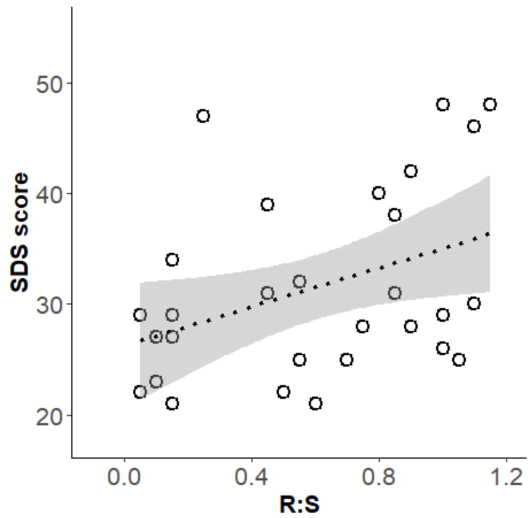


**Supplementary Figure 9. Depression and reward to shock ratio.**

Higher sensitivity to negative outcomes denoted by reward to shock ratios (R:S) was associated with higher levels of depressive symptoms (SDS score).

| **Fixed Effect** | **ß** | **SEM** | ***z*** | ***p*** | |
|---|---|---|---|---|---|
| (intercept) | 2.95 | 0.13 | 23.23 | <0.001 | *** |
| BO.Outcome | -0.17 | 0.02 | -9.59 | <0.001 | *** |
| BO.Uncertainty | 0.17 | 0.03 | 6.63 | 0.03 | *** |
| BO.Replay | 0.06 | 0.02 | 3.03 | <0.001 | ** |
| MO.Outcome | -0.07 | 0.02 | -3.99 | <0.001 | *** |
| MO.Uncertainty | 0.54 | 0.03 | 21.44 | <0.001 | *** |
| MO.Replay | 0.01 | 0.02 | 0.60 | 0.55 | |
| WO.Outcome | -0.25 | 0.02 | -14.42 | <0.001 | *** |
| WO.Uncertainty | -0.15 | 0.03 | -5.53 | 0.03 | *** |
| WO.Replay | -0.13 | 0.02 | -6.05 | <0.001 | *** |
| Choice Difficulty | 0.95 | 0.03 | 31.40 | <0.001 | *** |
| Gamble Difficulty Bias | 0.29 | 0.03 | 8.70 | <0.001 | *** |
| Sequence Check Score | -0.05 | 0.02 | -2.81 | 0.005 | ** |
| RT | -0.27 | 0.02 | -12.74 | <0.001 | *** |
| BO.Outcome: BO.Replay | -0.06 | 0.02 | -3.62 | <0.001 | *** |
| BO.Uncertainty: BO.Replay | -0.03 | 0.02 | -1.57 | 0.12 | |
| MO.Outcome: MO.Replay | -0.04 | 0.02 | -2.39 | 0.07 | * |
| MO.Uncertainty: MO.Replay | -0.006 | 0.02 | -0.28 | 0.78 | |
| WO.Outcome: WO.Replay | 0.04 | 0.02 | -3.62 | 0.03 | * |
| WO.Uncertainty: WO.Replay | 0.03 | 0.02 | -1.57 | 0.03 | * |

**Supplementary Table 1. Predicting rational choice with replay.**

Fixed effects from the regression model examining the effect of the replay levels of the best option (BO), mid option (MO) and worst option (WO) on rational choice. Shaded effects are the replay effects reported in **Supplementary Note 6**.

| Questionnaire | Sub-score | DA | OC | AW |
|---|---|---|---|---|
| Depression (SDS) | Affective | 0.326 | 0.396 | 0.007 |
| | Cognitive | **0.908** | -0.182 | 0.087 |
| | Somatic | **0.762** | 0.060 | 0.033 |
| Impulsivity (BIS-11) | Attention | **0.585** | 0.266 | -0.337 |
| | Cognitive instability | 0.051 | 0.262 | -0.155 |
| | Motor | -0.191 | **0.512** | 0.009 |
| | Perseverance | 0.497 | 0.175 | -0.172 |
| | Cognitive Complexity | 0.268 | 0.266 | -0.171 |
| | Self-control | 0.451 | 0.233 | -0.017 |
| OCD (OCI-R) | Hoarding | 0.008 | **0.690** | 0.306 |
| | Washing | 0.189 | 0.234 | -0.113 |
| | Obsessing | 0.181 | **0.671** | -0.006 |
| | Ordering | -0.056 | **0.893** | 0.007 |
| | Checking | -0.049 | **0.794** | -0.025 |
| | Neutralising | 0.318 | 0.301 | -0.207 |
| Trait anxiety (STAI-Y2) | Anxiety | 0.185 | 0.354 | **0.626** |
| | Depression | **0.723** | 0.057 | 0.310 |
| Obsessive beliefs (OBQ-20) | Thought control | 0.383 | -0.100 | 0.053 |
| | Responsibility | 0.227 | -0.152 | 0.126 |
| | Threat | 0.352 | -0.063 | 0.347 |
| | Perfectionism | 0.372 | 0.127 | 0.121 |
| Worry (WDQ-SF) | | 0.066 | 0.011 | **0.966** |

**Supplementary Table 2. Factor analysis of mental health questionnaires.**

Item loadings of questionnaire sub-scores unto factors spanning depressive-affect' (DA), 'obsessive-compulsive' (OC) and anxious-worry' (AW) dimensions. High positive loadings (>0.5) for each factor are in bold.

### 3. Supplementary Notes

#### Supplementary Note 1: Calibrating rewards versus shocks

Prior gambling-style decision tasks often use the loss of monetary reward as the negative outcome[8], and it is known that humans do not weigh reward gain and loss equally[9]. Here, we opted to use unpleasant electric shocks instead of monetary loss given its role as a primary punisher[10], that has individual differences in the valuation of money versus shocks and follow a grossly linear relationship between them[11]. We thus designed a reward to shock ratio calibration task in order to balance the gain of a positive outcome with that of a negative outcome, in order to transform the outcome values in the decision task in a participant-specific manner (**Supplementary Fig. 1A**). This ensured that the valuation of the positive (money) and negative (shocks) outcomes were equal across participants, and removes the bias of overweighing negative versus positive outcomes.

In the calibration paradigm, participants were shown various unit combinations of monetary reward and shocks (e.g., £0.5 and 1 shock) and were told to accept or reject the combination (Fig. S2A). If they accepted, the combination may be selected as an actual consequence (i.e., they will earn £0.5 as bonus, but will also receive 1 shock unit). The combinations were controlled by a two-down one-up staircase procedure, where the value of money decreased after two accepts and increased after one reject. We interleaved two staircases with n=30 trials (combinations) presented per staircase. Each staircase was initialised to £0.50 in ratio to 1 shock, had a target probability of 50% acceptance/rejection responses with a step size of +/-£0.10, with a £0.10 minimum (except 1 participant at £0.05) and £2 maximum. On every trial, the magnitude of reward and shocks were randomly scaled by 1-5, but the ratio between them remained as determined by the staircase. The mean of the final values from the two staircases was taken as the reward-shock ratio (R:S). This ratio, that ranged from 0.05-1.15 (M=0.60, SD=0.37), was then subsequently utilised in the main experimental task to transform the original reward value (**Supplementary Fig. 1B**).

Post-hoc, we observed the comparison of behaviour quantified by the R:S transformed values in the decision trials at face value (values on the screen; SV) versus the participants' internal valuation (IV, values used in the main analyses) of them. Using IV, participants overall made more rational choices ($t(29)=2.24$, 95%CI=[0.17 3.66], $p=0.03$), particularly gamble ones ($t(29)=4.46$,

95%CI=[3.00 8.09], p<0.001), than if performed via SV (**Supplementary Fig. 1C**). Participants with higher R:S ratios (require more money to balance 1 shock; more sensitive to shocks) were also revealed to be more risk-seeking (higher probability of choosing gamble over certain when both options were of equal EV (indifference point)) when using IV than SV values (**Supplementary Fig. 1D**). In other words, without considering the participants' interval valuation of the outcome values, participants may erroneously appear to be less rational or risk-seeking in their choices. Together these observations demonstrate the necessity of balancing to remove decision biases confounded by the imbalance of valuating positive versus aversive outcomes.

**Supplementary Note 2: Decision behaviour**

We saw that participants' choices were guided by the EV of the options. Thus, the EV of the gamble option was generally higher than the certain option for gamble choices (M=2.02, SD=0.48; t(29)=22.88, p<0.001, 95% CI=[-1.84 2.20] (against EV=0)), and vice versa when choosing the certain option (M=-2.20, SD=0.45; t(29)=-27.01, p<0.001, 95% CI=[-2.37 -2.03] (against EV=0)) (**Supplementary Fig. 2A**). Participants also made more gamble (M=59.16%, SD=8.97%) than certain choices (t(29)=5.59, p<0.001, 95% CI=[11.61 25.02]) as would be the case for a fully rational participant (t(29)=1.20, p=0.24, 95% CI=[-1.37 5.28]) (**Supplementary Fig. 2B**) given the experimental schedule (**Supplementary Note 3**).

To probe whether the distinct components—options' outcomes and probabilities—were additionally influencing the decision process, we used a multi-level logistic regression (controlling for reaction time and the trial block's path sequence memory score). We assessed the impact of each path's outcome value (Outcome) and transition probability (Probability) on Choice (Certain/Gamble) with the model:

Choice ~ Certain Path Outcome + (Gamble Path 1 Outcome × Gamble Path 1 Probability) + (Gamble Path 2 Outcome × Gamble Path 2 Probability) + RT + Seq Check + (1 | Subject)

We found that participants tended to choose the gamble option if the certain option had low outcome values (β=-2.33, SE=0.10, p<0.001) (**Supplementary Fig. 2C**), as per the EV analysis (EV in certain path = outcome). However, beyond that, we found that participants choose the gamble option more often if the gamble options had high outcome values (more likely path: β=1.51, SE=0.06, p<0.001; less likely path: (β=0.69, SE=0.05, p<0.001). While higher probabilities did not directly contribute to making more gambles (β=-0.06, SE=0.04, p=0.13), we found that it modulated the gamble options' outcomes to affect choice, such that the higher probability the greater the effect of outcome value on choice (more likely path: β=0.44, SE=0.05, p<0.001; less likely path: (β=0.36, SE=0.05, p<0.001).

**Supplementary Note 3: Decision trial schedules**

Participants were randomly assigned to one of five schedules that determined the decision trials' option outcome values (reward, neutral, shock; 1 to 5 in magnitude) and probabilities (50%/50%, 70%/30%, 90%/10% gamble) (**Supplementary Fig. 3**). We allowed the trials to have a variety of outcome type combinations, where one option path had a neutral outcome and the other two option paths ended up in both reward, both shock, or one reward and one shock (i.e., three outcome combination types: reward-neutral-shock (RNS), reward-neutral-reward (RNR) or shock-neutral-shock (SNS)).

Within these three outcome combination types, we also varied which outcome type was linked to a transition probability. For instance, every trial consisted of a certain path, a less risky gamble path, and a riskier gamble path (e.g., 100%, 70%, 30%; in order of certain-less risky-riskier). For an RNS combination type, the trial could be reward-neutral-shock (RNS), reward-shock-neutral (RSN), neutral-reward-shock (NRS), neutral-shock-reward (NSR), shock-reward-neutral (SRN), shock-neutral-reward (SNR). For a RNR combination type, the trial could be reward-neutral-reward (RNR), reward-reward-neutral (RRN), neutral-reward-reward (NRR). For a SNS combination type, the trial could be shock-neutral-shock (SNS), shock-shock-neutral (SSN), neutral-shock-shock (NSS).

Therefore, there were altogether 12 different outcome-probability combination trial types. The five schedules were designed to ensure variability of options' outcome magnitudes whilst keeping specific numbers of outcome-probability combination trial types and an equal number of certain/gamble choices for a fully rational participant.

### Supplementary Note 4: Trial-to-trial independence of decisions

We conducted several control analyses to ensure that choices in the task were void of sequential effects from prior decisions. We found that the previous trial's outcome, prediction error (estimated as the difference between outcome and chosen option expected value), difficulty (i.e., absolute EV difference between options), choice type (certain or gamble), choice rationality (rational or irrational choice), or any interactions between these variables, did not predict the current trial's choice type ($ps>0.19$) nor choice rationality ($ps>0.18$). Thus, there were no significant behavioural influence of the decision trials unto subsequent ones.

**Supplementary Note 5: Inconsistent backward replay effects during deliberation time**

Given that significant backward sequenceness of the option paths were present during deliberation, we assessed whether it was linked to the path's outcome value and uncertainty, and whether this was associated with choice rationality.

We first used a regression model containing outcome value or uncertainty (plus covariates; see **Methods**) to predict backward replay level of the option path. While there was no significant effect of uncertainty on replay ($\beta$=0.0004, SE=0.0002, p=0.11), we saw that option paths with more positive outcomes had stronger backward sequenceness ($\beta$=0.0005, SE=0.0002, p=0.04). The latter effect seemed to be driven primarily by neutral outcomes where only neutral paths ($\beta$=-0.001, SE=0.0006, p=0.02), and not shock paths ($\beta$=-0.0008, SE=0.0006, p=0.14), were replayed more than rewarding paths. Moreover, the outcome on replay effect did not remain significant when we included both outcome value and uncertainty, with an interaction effect of choice rationality, into the same model. Both uncertainty ($\beta$=0.0006, SE=0.0006, p=0.32) and outcome ($\beta$=0.0005, SE=0.0006, p=0.42) of the path were not linked to its backward replay strength, and neither were their interactions with rational choice (ps>0.66). Given that the relationship of outcome and backward replay was not robust, we opted to refrain from concluding it as a true effect.

**Supplementary Note 6: Predicting rational choice with replay**

We examined uncertainty, outcome, and choice rationality effects on replay in the main text. However, this precluded a direct prediction of choice as the regression model would require all three paths to be included as fixed effects. For completeness, here we asked whether replay of individual choice options was predictive of whether a participant made an irrational choice. We ranked each of the three choice option paths within each trial by its objective desirability via its expected value (EV; highest EV option = best option (BO), mid EV option = mid option (MO), lowest EV option = worst option (WO)). We then assessed if replay of each option path contributed to the making of rational choice with a multi-level logistic regression that included the option paths' outcome values (Outcome) and uncertainty (Uncertainty) levels, as well as the choice and gamble difficulty biases (absolute and signed option EV difference), the trial block's path sequence memory score (Seq Check) and reaction time (RT). The model was:

Rational Choice ~ (BO.Outcome + BO.Uncertainty) × BO.Replay + (MO.Outcome + MO.Uncertainty) × MO.Replay + (WO.Outcome + WO.Uncertainty) × WO.Replay + Choice Difficulty + Gamble Difficulty Bias + Seq Check + RT + (1 | Subject / Lag)

Behaviourally, increasing outcome values for the best option and decreasing uncertainty contributed to rational choice selection (**Supplementary Table 1**). For the mid and worst options, decreasing outcome values and increasing uncertainty aided rational choice, likely due to making the best option more obvious.

Neural replay was also indeed predictive of rational choice, over and above the behavioural effects (**Supplementary Table 1**). If participants replayed the objectively best choice option more (highest EV), then this predicted rational choice ($\beta$=0.06, SE=0.02, $p$=0.002), whilst replaying the worst path (lowest EV) led to more irrational choice behaviour ($\beta$=-0.13, SE=0.02, $p$<0.001). There was no significant relationship of the overall sequenceness strength of a mid-value path on rational choice ($\beta$=0.01, SE=0.02, $p$=0.55). These results demonstrate that what participants replay is predictive of whether they make good or poor choices, and where a replay of bad options is linked to suboptimal decisions.

**Supplementary Note 7: Cross validation of choice prediction without and with replay**

We further tested how much replay contributed to choice, over and above behaviour. We thus used a 5-fold cross validation procedure, with 100 repetitions, to compare model fits for a model predicting choice with just behaviour, with behaviour and replay, or with behaviour and replay that had been randomly shuffled (i.e., made into noise).

The behaviour only model was:

Gamble Choice ~ CertainPathOutcome + GamblePath1Outcome × GamblePath1Probability + GamblePath2Outcome × GamblePath2Probability + RT+ Seq Check + Choice Difficulty + Gamble Difficulty Bias + (1| Subject / Lag)

The models including replay was:

Gamble Choice ~ CertainPathOutcome × CertainPathReplay + (GamblePath1Outcome × GamblePath1Probability) × GamblePath1Replay + (GamblePath2Outcome × GamblePath2Probability) × GamblePath2Replay + RT + Seq Check + Choice Difficulty + Gamble Difficulty Bias + (1| Subject / Lag)

In both cases, we ensured that rank deficiency due to the collinearity of GamblePath1Probability and GamblePath2Probability (contingency of gamble paths always add up to 100%) was dealt with by replacing affected regressors (GamblePath1Probability = GamblePath2Probability). To account for the nested random effect of Subject/Lag, we clustered the partitions for each fold at the participant and lag level.

Over 100 repetitions, we found that the inclusion of replay decreased the root mean square error (RMSE) (vs. behaviour: t(99)=-48.88, 95% CI=[-0.00048 -0.00044], p<0.001; vs. permuted replay: t(99)=- 38.67, 95% CI= -0.00053 -0.00048], p<0.001) and increased the adjusted $R^2$ (vs. behaviour: t(99)=369.92, 95% CI=[0.00449 0.00453], p<0.001; vs. permuted replay: t(99)=241.26, 95% CI=[0.0042 0.0043], p<0.001) and likelihood (vs. behaviour: t(99)=146.78, 95% CI=[0.099 0.10], p<0.001; vs. permuted replay: t(99)=97.34, 95% CI=[0.097 0.10], p<0.001) in predicting choice when compared to both models of just behaviour or with permuted replay (**Supplementary Figure 4**).

**Supplementary Note 8: Choice difficulty effects on replay strength**

How much we deliberate or reflect on decisions is often dependent on how difficult the choice is. In the decision trials, choice difficulty was represented as both (i) the signed expected value difference between gamble minus certain options (where a positive value denoted an easier gamble choice) and (ii) the absolute expected value difference between gamble versus certain options (where a positive value denoted an easier overall choice).

We observed that during deliberation phase, only overall lower choice difficulty was linked to decreased backward replay strength ($\beta$=-0.004, SE=0.001, p<0.001), which was driven by lower replay in easier trials prior to irrational choices (interaction effect: $\beta$=0.004, SE=0.001, p<0.001) (**Supplementary Fig. 5A**). For post-choice replay, we found that lower gamble choice difficulty was linked to decreased forward replay strength ($\beta$=-0.01, SE=0.004, p=0.007), which was driven by lower replay in easier gamble choice trials after irrational choices (interaction effect: $\beta$=0.009, SE=0.004, p=0.02) (**Supplementary Fig. 5B**). All other effects were non-significant. We included these two variables as co-variates in our analyses to control for choice difficulty effects on replay and choice rationality.

**Supplementary Note 9: Quantifying counterfactual prediction error after choice**

In quantifying counterfactual prediction error, we considered three definitions:

Def 1: PE = Each Path Outcome –Trial's Experienced Path Outcome

Def 2: PE = Each Path EV – Trial's Experienced Path EV

Def 3: PE = Each Path Outcome –Trial's Chosen Option EV

Def 1 analyses are reported in the main text.

For Def 2, PE has the same meaning to Def 1. Options which the participant missed out from their choice (inexperienced paths) would have a positive PE if they led to a better outcome than the one experienced (regret) and a negative PE if that path would have led to a worse outcome than experienced (relief). Associations of this counterfactual PE and its interaction with choice rationality predicting backward sequenceness strength is the same as those of Def 1. Paths with negative PEs had higher backward replay ($\beta$=-0.009, SE=0.002, $p<0.001$) overall (**Supplementary Fig. 6A**). The significant interaction effect of PE with choice rationality was present ($\beta$=0.01, SE=0.002, $p<0.001$), where paths with lower PEs (i.e. worse alternative paths than the experienced path) had stronger backward sequenceness after irrational choice ($\beta$= -0.009, SE=0.002, $p<0.001$), but had weaker backward sequenceness after rational choices ($\beta$=0.002, SE=0.0007, $p$=0.005). These results can be interpreted in the same idea that participants engaged in a form of relief processing (i.e. did not get worst option) after irrational choices, but a disappointment (i.e. did not get the best option) after making rational choice. This counterfactual PE was also a better model fit than outcome value (PE vs outcome model: $\Delta$AIC=-21.61).

For Def 3, PE is constituted by the difference between the EV of the trial's chosen option (certain or gamble) versus each of the three outcome paths participants could receive. In other words, positive PE values reflected option paths which were better than expected, whilst negative PE values reflected option paths which were worse than expected. As for its link to backward replay, we likewise saw that backward sequenceness strength increased for option paths where PE was more negative (β=-0.005, SE=0.002, p=0.004), suggesting that worse than expected option paths were linked to stronger backward sequenceness overall (**Supplementary Fig. 6B**). We also saw a significant interaction effect of PE with choice rationality (β=0.006, SE=0.002, p<0.001), In separate models of just rational or irrational trials, paths with lower PEs (i.e. worse outcomes than expected) had stronger backward sequenceness after irrational choice (β=-0.005, SE=0.002, p=0.005), but had weaker backward sequenceness after rational choices (β=0.002, SE=0.0007 p=0.03). This meant that backward sequenceness increased for worse outcomes than expected after irrational choice, but that after rational choice even better outcomes than expected were more prominently replayed. Again, path sequenceness strength was better explained by this counterfactual PE value than outcome value alone (PE vs outcome model: ΔAIC=-7.11). However, whether these effects can be classified as regret/relief processes are less clear, given that PE≠0 included paths that were also experienced.

**Supplementary Note 10: Replay patterns in mental illness**

We investigated individual differences in mental health symptoms and their relationship to replay, decision-making, and irrational choice. We assessed mental health traits using self-report questionnaires encompassing anxiety and also common co-occurring and overlapping symptoms like worry, depression, obsessive-compulsive traits, and impulsivity. To reduce the collinearity of the (sub)scores (r=-0.24 to 0.80), we conducted a factor analysis resulting in three latent factors (r=0.20 to 0.41) encompassing anxious-worry (AW), obsessive-compulsive (OC) and depressive-affect (DA) dimensions. We then tested the impact of these symptoms on anticipatory replay (by including the mental health factors in above models predicting replay during deliberation; correcting for multiple comparisons with Bonferroni correction for the three factors) using the model:

$$\text{Replay} \sim (\text{Outcome} + \text{Uncertainty} + \text{Choice Difficulty} + \text{Gamble Difficulty Bias}) \times \text{Rationality} \times (\text{AW} + \text{OC} + \text{DA}) + \text{RT} + \text{Seq Check} + (1 \mid \text{Subject} / \text{Lag})$$

During deliberation time, we found significant effects (interactions) of obsessive-compulsion and anxious-worry on outcome value and choice rationality. Participants with high obsessive-compulsive levels seemed to exhibit an over-focus on aversive outcomes, as they had stronger forward sequenceness for more aversive option paths ($\beta=-0.003$, SE=0.0008, $p<0.001$, corr.) (**Supplementary Fig. 8A**). Interestingly, this was particularly pronounced before making irrational choices (interaction: $\beta=0.003$, SE=0.0009, $p=0.003$, corr.). The latter effect was similarly captured by a model examining only irrational trials—obsessive-compulsive scores were linked to stronger replay for aversive paths in these trials ($\beta=-0.003$, SE=0.0009, $p=0.001$, corr.) (**Supplementary Fig. 8A**). Whilst this biased replay in obsessive-compulsive subjects did not translate into making more irrational choices per se ($\beta=-0.07$, SE=0.09, $p=0.40$, uncorr.), it does align well with the psychological concept of OCD where an over focus on potential negative outcomes is believed to drive compulsions[12,13]

In contrast, people with high anxious-worry symptoms showed a decreased replay for good outcomes, particularly before making rational choices (**Supplementary Fig. 8B**; rational choices only: $\beta=-0.002$, SE=0.0004, $p<0.001$, corr.; interaction with rationality across all trials: $\beta=-0.003$,

SE=0.001, p=0.02, corr.). This meant that people with high anxious-worry symptoms focused disproportionately less on rewarding outcomes, aligned with worry ruminations which are often utilised to problem solve or self-blame[14].

Interestingly, these effects were present even after experimentally controlling for biased perceptions of aversive outcomes (i.e., electric shocks in our experiment). This was done by weighing negative outcomes more than positive ones with a reward to shock ratio (R:S) that represented an individuals' valuation of 1 unit of shock in terms of monetary reward (see **Supplementary Note 1**). Higher R:S individuals were those who required more money to receive 1 unit of shock, which meant that they were more sensitive to the aversive stimuli. We found that individuals who had elevated levels of depressive symptoms also had higher R:S values (r=0.39, p=0.03) (**Supplementary Fig. 9**), in line with the prior literature[15].
